# Mutually inclusive mechanisms of drought-induced tree mortality

**DOI:** 10.1101/2020.12.17.423038

**Authors:** Peter Hajek, Roman M. Link, Charles Nock, Jürgen Bauhus, Tobias Gebauer, Arthur Gessler, Kyle Kovach, Christian Messier, Alain Paquette, Matthias Saurer, Michael Scherer-Lorenzen, Laura Rose, Bernhard Schuldt

## Abstract

An extreme summer drought caused unprecedented tree dieback across Central Europe in 2018, highlighting the need for a better mechanistic understanding of drought-induced tree mortality. While numerous physiological risk factors have been identified, the principal mechanisms, hydraulic failure and carbon starvation, are still debated. We studied 9,435 trees from 12 temperate species planted in a diversity experiment in 2013 to assess how hydraulic traits, carbon dynamics, pest infestation, tree height and neighbourhood competition influence individual mortality risk. We observed a reduced mortality risk for trees with wider hydraulic safety margins, while a rising sugar fraction of the non-structural carbohydrate pool and bark beetle infestation were associated with higher risk. Taller trees had a lower mortality risk. The sign and magnitude of neighbourhood effects on mortality risk depended on the species-identity of the involved trees, with most species having beneficial and some having deleterious effects on their neighbours. While severe tissue dehydration causing hydraulic failure precedes drought-induced tree mortality, we show that the probability of this event depends on a series of mutually inclusive processes including pest infestation and starch depletion for osmotic adjustment, and is modulated by the size and species identity of a tree and its neighbours.

## Introduction

Worldwide, forests are exposed to rises in temperature, atmospheric vapour pressure deficit and intensity and frequency of severe drought events (Dai 2013; Yuan et al. 2019; Zhou et al. 2019). As a consequence, large-scale tree mortality events are documented for all forest biomes (Allen et al. 2015; Hartmann et al. 2018; Senf et al. 2018; Schuldt et al. 2020). Challenges in forecasting the impacts of drought-related tree mortality based on the structure and composition of forests highlight the need for a robust mechanistic understanding of the processes involved (Choat et al. 2018; Brodribb et al. 2020).

Two interrelated physiological mechanisms are central for drought-induced tree mortality: hydraulic failure, which is the partial or complete loss of xylem functionality due to embolism formation, and carbon starvation, the depletion of non-structural carbohydrates (NSC) due to negative drought impacts on photosynthesis (McDowell et al. 2008). While there is clear evidence that hydraulic failure is a universal component of the processes preceding tree death under drought (Rowland et al. 2015; Anderegg et al. 2016; Adams et al. 2017; Correia et al. 2019; Powers et al. 2020; Li et al. 2020), the role of NSC reserves in these processes is still debated (Adams et al. 2017; Blackman et al. 2019; Kannenberg & Phillips 2020; O’Brien et al. 2020). Both processes interact with antagonistic biotic agent demographics (McDowell et al. 2008, 2011), as drought effects on the water and carbon balance may additionally predispose trees to pathogen or insect attacks (Raffa et al. 2008; Sturrock et al. 2011; Hart et al. 2014; Huang et al. 2020), exacerbating climate-driven forest mortality (McDowell et al. 2013, 2020; Seidl et al. 2017).

Many risk factors have been associated with drought-induced mortality (O’Brien et al. 2017). Specifically, increasing tree size is often found to increase drought vulnerability (Peng et al. 2011; Bennett et al. 2015; O’Brien et al. 2017; Stovall et al. 2019). Further, water shortage may intensify interspecific tree competition depending on species-specific water use behaviour and above- and below-ground niche partitioning (Ammer 2019; Grossiord 2019), thus affecting individual mortality risk via variation in local neighbourhood composition (Vitali et al. 2018; Fichtner et al. 2017, 2020).

In the recent past, several distinct mechanisms of drought-induced mortality have been proposed for a variety of tree species in different biomes (Adams et al. 2017; O’Brien et al. 2017; Choat et al. 2018; Brodribb et al. 2020). Mechanistic insight advancing future process-based models is most likely to result from research that accounts for the multitude of simultaneous processes at play during tree death under drought (McDowell et al. 2019). Consequently, field-based studies that jointly analyse multiple potential causes of drought-induced tree mortality across a range of functionally different tree species are urgently needed. In 2018, Central Europe experienced an extreme summer drought, followed by a non-typical dry winter and spring (Hari et al. 2020). This global-change type drought event resulted in the highest average growing season temperature and vapour pressure deficit ever recorded (3.3°C and 3.2 hPa above the long-term average for the reference period from 1961-1990, respectively) and caused unprecedented drought-induced tree mortality in many species (Schuldt et al. 2020). We used observations from 9,435 trees belonging to six angiosperm and six coniferous species from Europe and North America grown in a young tree diversity and nutrient enrichment experiment established in 2013 in Southern Germany (Wein et al. 2016) to try to untangle the complexity of possible mechanisms explaining drought-induced mortality in trees. Based on a Bayesian hierarchical modelling approach, we predicted the individual mortality risk following the drought using data on hydraulic traits, carbon dynamics, pest infestation, tree height and interactions with neighbouring trees. We ask whether (i) a tree species’ susceptibility to drought-induced mortality depends on both hydraulic traits and NSC dynamics, and whether (ii) the probability of dying on the individual level is modulated by tree height and the influence of biotic (pest infestation and interaction with neighbour trees) and abiotic (fertilisation and environmental) factors.

## Results and discussion

### Cross-species patterns of drought-induced tree mortality rates

In response to the anomalous severe drought of the year 2018 and the prolonged dry conditions that followed throughout the winter and spring of 2019, 34% of the 9,435 trees present in our experimental plantation died. Tree mortality increased dramatically compared to a total loss of 9% of trees between 2013 and early 2018 (see Methods). Mortality rates varied strongly across the 12 studied species, ranging from 0.6% for *A. saccharum* to 79.5% for *L. laricina* (Table 1, Fig. 1). The average probability of dying predicted by our model closely reflected the specieslevel mortality rates observed in the field (Fig. 1). Our model reached full convergence (Table S2.1) and explained 51.8% of the variance in the observed individual mortality (Fig. 2). Mortality rates were credibly elevated over the average mortality rate (i.e. their 95% highest posterior density interval (HDI) excluded the overall average) for *Larix* spp., *Betula* spp. and *P. abies*, while they were much lower specifically for *Acer* spp. and *Q. rubra* (credibly below 10% in all three cases).

**Fig. 1.**
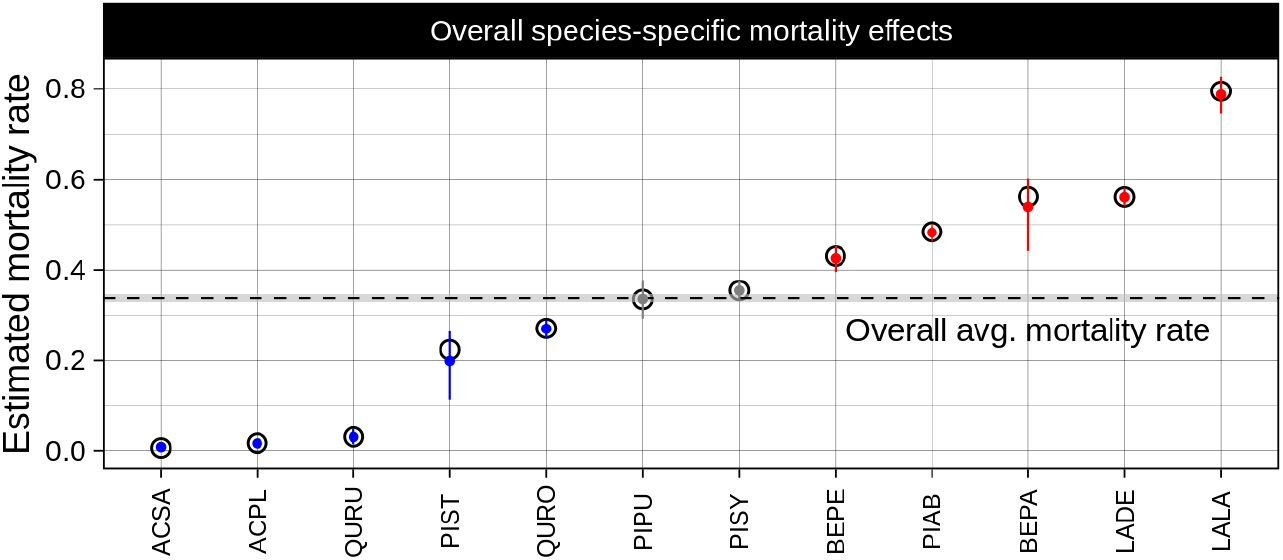
Average estimated mortality rates for each species ordered by magnitude (posterior mean ± 95% highest posterior density interval (HDI)). Dashed line with grey ribbon: overall predicted average ± 95% HDI. Red: risk credibly elevated over average level; blue: risk credibly below average; grey: not credibly different from average. For species acronyms, see Table 1. Black circles: observed average mortality rates (mismatch for *P. strobus, B. papyrifera* and *L. laricina* results from misclassification).

**Fig. 2.**
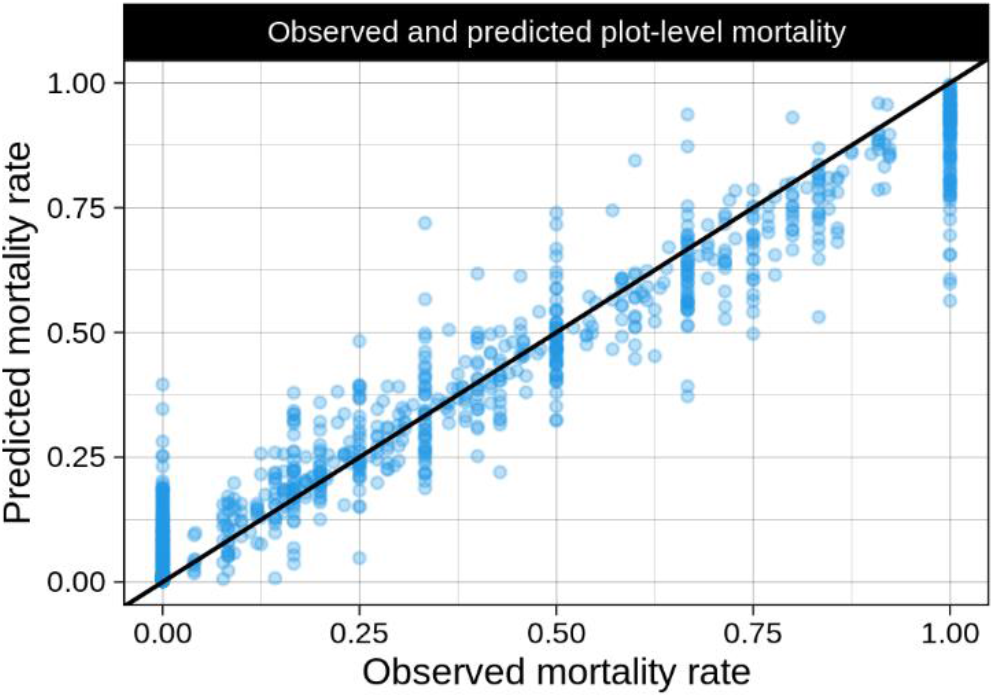
Posterior mean species-wise plot average mortality rate vs. observed proportion of dead trees per species and plot (explained variance in individual tree mortality: 51.8%).

**Table 1.** Properties of the 12 studied species. Given are the acronyms, species names, origin (EU: Europe; NA: North America), group (angio- or gymnosperm), pre-drought height, stem diameter at 10 cm height (Dia.), xylem pressure at 50% and 88% loss of conductivity (*P*_50_ and *P*_88_), midday water potential (*Ψ*_md_), change in leaf NSC sugar fraction over the growing period (Δsugar), proportion of trees with visible beetle damage (% of central trees), proportion of trees with neighbours infested by beetles (% of central trees), and observed mortality (% of central trees). Given values are mean ± SE. *P*_50_ and *P*_88_-values for the two *Quercus* species were taken from Lobo et al. (2018).

| Acronym | Species | Origin | Group | Height (m) | Dia. (mm) | $P_{50}$ (MPa) | $P_{88}$ (MPa) | $\Psi_{md}$ (MPa) | HSM (MPa) | $\Delta$ sugar (%) | B. (%) | N.w.B. (%) | Mort<sub>obs</sub> (%) |
| --- | --- | --- | --- | --- | --- | --- | --- | --- | --- | --- | --- | --- | --- |
| ACPL | *Acer platanoides* | EU | Angio | 4.30 ± 0.04 | 33.0 ± 0.4 | -4.56 ± 0.16 | -5.27 ± 0.12 | -2.50 ± 0.05 | 2.77 ± 0.06 | -21.47 ± 7.86 | 0.0 | 8.8 | 1.6 |
| ACSA | *Acer saccharum* | NA | Angio | 4.72 ± 0.07 | 30.1 ± 0.6 | -4.22 ± 0.11 | -4.61 ± 0.13 | -2.83 ± 0.25 | 1.78 ± 0.25 | -24.31 ± 7.48 | 0.0 | 3.8 | 0.6 |
| BEPA | *Betula papyrifera* | NA | Angio | 4.92 ± 0.05 | 43.0 ± 0.7 | -2.29 ± 0.03 | -2.42 ± 0.04 | -1.66 ± 0.17 | 0.63 ± 0.17 | 3.56 ± 2.14 | 0.0 | 0.3 | 56.1 |
| BEPE | *Betula pendula* | EU | Angio | 5.26 ± 0.03 | 50.0 ± 0.5 | -2.33 ± 0.04 | -2.47 ± 0.05 | -1.30 ± 0.05 | 1.02 ± 0.05 | 16.16 ± 10.4 | 0.2 | 2.4 | 43.0 |
| LADE | *Larix decidua* | EU | Gymno | 3.75 ± 0.04 | 39.8 ± 0.5 | -3.63 ± 0.07 | -4.05 ± 0.10 | -1.98 ± 0.09 | 1.65 ± 0.09 | 17.00 ± 3.08 | 15.8 | 29.1 | 56.1 |
| LALA | *Larix laricina* | NA | Gymno | 3.18 ± 0.07 | 27.7 ± 0.8 | -3.44 ± 0.05 | -3.85 ± 0.09 | -2.01 ± 0.05 | 1.43 ± 0.05 | -3.43 ± 7.75 | 0.0 | 3.6 | 79.5 |
| PIAB | *Picea abies* | EU | Gymno | 2.06 ± 0.02 | 27.1 ± 0.2 | -3.77 ± 0.10 | -4.53 ± 0.17 | -2.11 ± 0.31 | 1.67 ± 0.31 | -19.82 ± 5.37 | 12.8 | 21.4 | 48.4 |
| PIPU | *Picea pungens* | NA | Gymno | 1.16 ± 0.02 | 22.7 ± 0.4 | -4.04 ± 0.07 | -4.84 ± 0.11 | -1.96 ± 0.04 | 2.08 ± 0.04 | -10.06 ± 5.63 | 1.4 | 9.8 | 33.4 |
| PIST | *Pinus strobus* | NA | Gymno | 2.52 ± 0.05 | 33.0 ± 0.6 | -2.97 ± 0.03 | -3.26 ± 0.04 | -1.76 ± 0.05 | 1.21 ± 0.05 | -33.19 ± 1.75 | 0.0 | 2.8 | 22.3 |
| PISY | *Pinus sylvestris* | EU | Gymno | 2.70 ± 0.03 | 36.8 ± 0.5 | -3.10 ± 0.05 | -3.57 ± 0.04 | -1.84 ± 0.12 | 1.26 ± 0.12 | -1.48 ± 4.71 | 1.4 | 11.2 | 35.4 |
| QURO | *Quercus robur* | EU | Angio | 2.21 ± 0.03 | 18.5 ± 0.3 | -4.74 ± 0.09 | -5.66 ± 0.09 | -3.00 ± 0.03 | 2.66 ± 0.10 | 0.00 ± 0.00 | 0.1 | 5.9 | 27.1 |
| QURU | *Quercus rubra* | NA | Angio | 3.54 ± 0.08 | 24.3 ± 0.6 | -4.43 ± 0.25 | -6.72 ± 0.29 | -3.07 ± 0.09 | 3.65 ± 0.30 | -14.05 ± 11.2 | 0.0 | 6.6 | 3.0 |

### Hydraulic safety margins and non-structural carbohydrates

Both hydraulic traits and carbohydrate dynamics differed considerably among species, with observed hydraulic safety margins (HSM) ranging from 0.63 to 3.65 MPa, and changes in the fraction of soluble sugars (glucose, fructose and sucrose) to total leaf NSC (Δ_sugar_) over the growing season ranging from −33.2 to +17.0% (Table 1). An increase in relative leaf sugar content is indicative of starch depletion and higher need of soluble sugars for osmotic adjustment (Hsiao 1976; Thalmann & Santelia 2017). We detected credible effects (i.e. 95% HDI excluding zero) of both HSM and Δ_sugar_ on the mortality risk, with species with a wider HSM being at lower risk, and species with a more pronounced increase in relative leaf sugar content dying at higher rates (Fig. 3, Fig. 4, Table S2.1). This suggests that drought-induced mortality in trees is jointly influenced by both hydraulic failure and carbon starvation (McDowell et al. 2011).

**Fig. 3.**
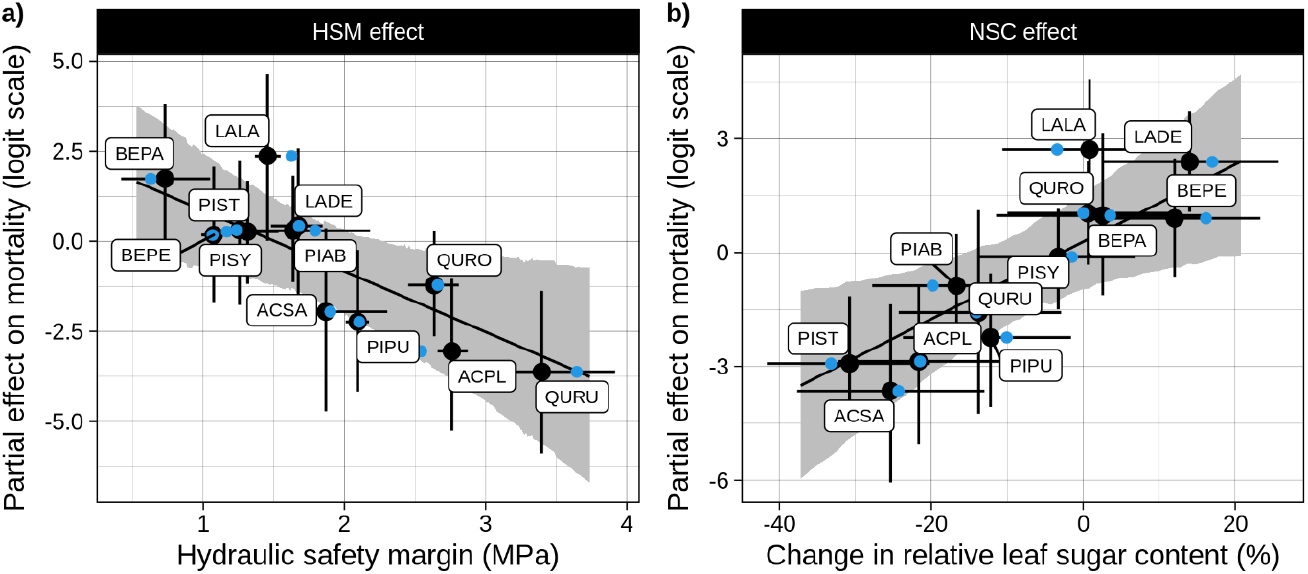
Partial effects for the species level effects of a) hydraulic safety margins and b) change in relative leaf sugar content (indicative of starch depletion) on the mortality risk. Black line with grey bands: posterior mean ± 95% HDI. For species acronyms, see Table 1. Blue points: average observed values; black points: estimates ± 95% HDI.

**Fig. 4.**
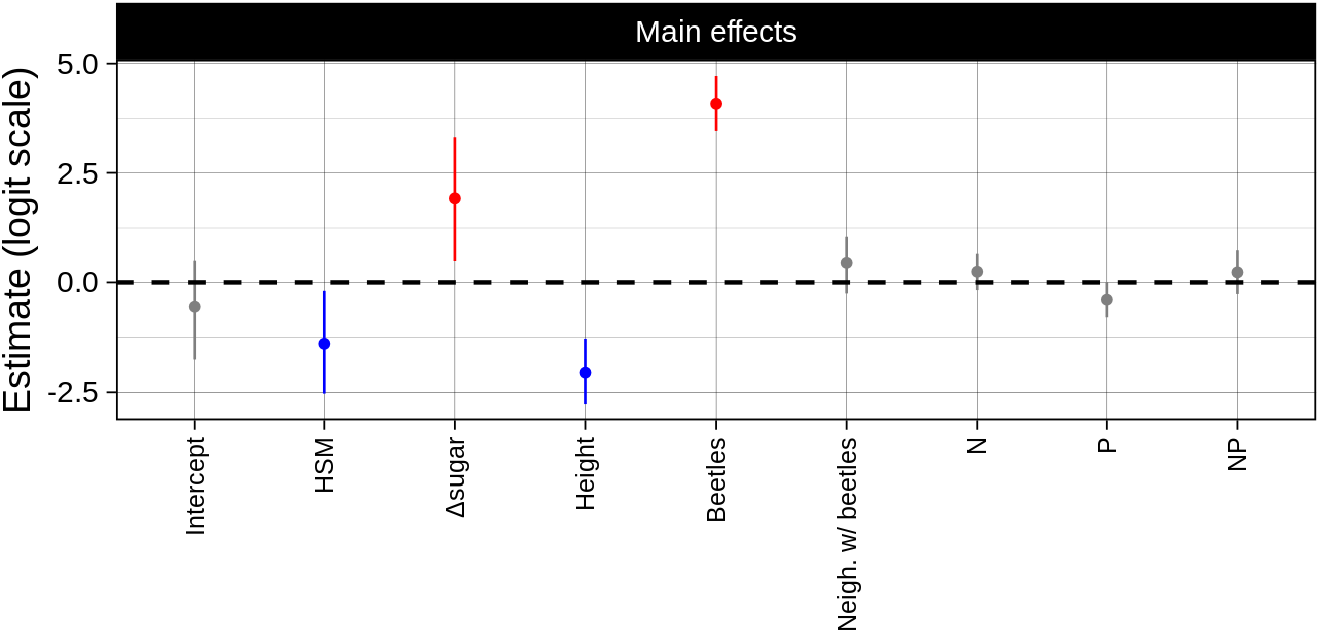
Estimated main effects on mortality on the logit scale (posterior mean ±95% HDI). Blue: credibly reduced risk; red: credibly elevated risk; grey: no credible effect on mortality. HSM: hydraulic safety margins. Compare Fig. S2.2 for species-specific effects.

The lower mortality we observed in species with wider HSMs is in line with reported crossspecies patterns in all forested biomes (Anderegg et al. 2016) and supports the hypothesis that hydraulic traits capture proximate mechanisms determining tree death (Adams et al. 2017). After the loss of hydraulic connectivity at the soil-root-interface during prolonged drought (Carminati & Javaux 2020), the amount of internally stored water and its loss via leaky stomata, the cuticle and the bark determine the timing of lethal tissue dehydration (Blackman et al. 2016; Choat et al. 2018; Duursma et al. 2019). Reaching critical desiccation thresholds may thus mark the final step in the dying process of trees rather than being the cause of death.

In contrast to hydraulic failure, the diverse physiological responses affecting NSC reserves at the time of death indicate that carbon starvation is not a universal phenomenon of tree death during drought events (Hartmann et al. 2013; Adams et al. 2017; Dai et al. 2018; O’Brien et al. 2020). However, partial canopy dieback and/or the loss of xylem functionality will result in a reduction of leaf area, which in turn affects whole-plant net photosynthesis and carbon allocation in post-drought years. Recent evidence suggests that embolism repair under tension, i.e. restoration of xylem functionality concurrent with transpiration, is either a rare phenomenon or even completely absent in many tree species (Charrier et al. 2016; Choat et al. 2019; Duan et al. 2019; Rehschuh et al. 2020; but see Klein et al. 2018; McDowell et al. 2019). This might explain commonly observed legacy effects of reduced vigour and increased mortality rates up to several decades after a drought event (Cailleret et al. 2017; Timofeeva et al. 2017; Trugman et al. 2018), as well as different patterns of post-drought recovery (Li et al. 2020). The mechanisms governing drought-induced tree mortality and recovery might therefore be closely linked to carbon dynamics (Trugman et al. 2018). Noteworthily, *Larix* spp. and *Betula* spp., two of the most strongly drought-affected genera, showed signs of leaf browning and defoliation already in early July 2018. While leaf shedding can mitigate drought effects on a plants’ water balance via reduction of the transpiring surface (Blackman et al. 2019), the high Δ_sugar_ in these genera (Table 1) indicates that the reduced carbon gain caused by this strategy may have contributed to their high mortality rates.

Both access to carbohydrate reserves and their utilization rate have been reported to be controlled by water availability (Sevanto et al. 2014), providing evidence that carbohydrate use is controlled by the functionality of the water transport system (Atkin & Macherel 2009) rather than by photosynthetic carbon gain. This is consistent with a hydraulic constraint on NSC consumption (McDowell & Sevanto 2010; Sala et al. 2010; McDowell et al. 2011). The role of carbohydrates in osmotic adjustment constitutes a central link between carbon metabolism and hydraulic functioning (Pantin et al. 2013; Merchant 2014). In the absence of drought, experimental manipulation of NSC storage can affect osmotic regulation and shift the turgor loss point towards substantially less negative water potentials (Sapes et al. 2020). This is consistent with a recent meta-analysis providing evidence for a decrease in starch and rise in soluble sugars over the course of extreme drought events (He et al. 2020). The observed decrease of the relative starch fraction in the more drought-affected species is a strong indication for starch-to-sugar conversion to meet the trees’ metabolic and osmoregulatory demands during drought. The use of stored starch to produce osmotically active sugars indicates that the supply with recent assimilates was not sufficient for osmotic regulation. For example, extreme drought led to a strong conversion of starch to sugars in needles and other plant tissues in Scots pine before a mortality event, while the starch-to-sugar ratios did not change during a milder and non-lethal drought (Schönbeck et al. 2020). The use of local storage for osmotic regulation thus seems to be an indicator of carbon starvation and does ensure survival under extreme drought. If plants indeed increase the concentrations of soluble sugars to maintain turgor under declining water potentials, the NSC depletion often seen in dying trees (Adams et al. 2017) is likely to act as an underlying cause of hydraulic failure rather than a separate process. This may explain the divergent explanations of the proximate causes of plant death under drought.

### Tree size effects

In addition to the species-level effects of HSM and NSC dynamics, our model identified a positive within-species effect of tree height on the survival of all species except for *Q*. *rubra*, though the magnitude of that height effect varied strongly among species (Fig. 4; Fig. S2.2).

This finding is in contrast to mounting evidence that larger trees tend to be more susceptible to drought-induced tree mortality across biomes (Phillips et al. 2010; Lindenmayer et al. 2012; Bennett et al. 2015; Grote et al. 2016; Olson et al. 2018; Stovall et al. 2019, 2020), although some authors report opposite patterns (Van Mantgem et al. 2009; Peng et al. 2011). This apparent contradiction might result because studies reporting higher mortality for the largest trees usually focus on adult trees in mature forest stands, where crown exposure (Stovall et al. 2019) and path length-related constraints on water transport (Ryan & Yoder 1997; West et al. 1999) are more relevant. Our analysis, in turn, was based on densely spaced and young trees (planted in 2013 at the age of 1-3 years) with limited variation in tree height (Table 1). Under these circumstances, negative density-dependent drivers of mortality are likely to dominate (Crouchet et al. 2019), disproportionally affecting smaller, competitively supressed tree individuals via self- and alien-thinning (Kohyama 1994, Pretzsch & Forrester 2017). A potential explanation for the competitive advantage of larger trees in the experiment is their extensive root system and better access to deep soil water, particularly during periods of water shortage (Ledo et al. 2018).

### Bark beetle effects on conifers

Individuals diagnosed with bark beetle infestation in 2018 had a much higher probability of dying (Fig. 4), consistent with a drought-driven predisposition for biotic attacks (McDowell et al. 2011). The prevalence of bark beetles differed greatly between species, with strong outbreaks only for *P. abies* and *L. decidua* (Table 1). Most likely, a decline in resin exudation and thus defence capability caused by the reduction of relative tissue water content made *P. abies* vulnerable to insect infestation (Netherer et al. 2015). Despite the low overall prevalence, the infested individuals among the native conifers *P. abies*, *L. decidua* and *P. sylvestris* died with high probability. As bark beetles cause physical damage to the phloem and xylem that is likely to affect both carbon metabolism and xylem water transport, their role for plant death cannot be understood without acknowledging its link to other processes driving drought-induced mortality (McDowell et al. 2008, 2011). Since both healthy and infested trees died during the drought event while a number of infested trees were able to survive, biotic attack was unlikely to be the main driver of mortality in the affected species. Therefore, pest infestations should rather be interpreted as an additional stressor amplifying the risk of death by other causes.

The presence of bark beetles in neighbouring trees had no overall effect on mortality (Fig. 4), and on species level only affected *P. abies*, the native species that suffered the strongest beetle infestation (Fig. S2.2). While neighbourhood effects can profoundly impact the predisposition to insect herbivory (Castagneyrol et al. 2018), the low influence in our study may be related to the timing of the assessment of pest infestation in late 2018, when the drought and associated insect outbreaks hat likely already run its course and further spread was unlikely.

### Interactions with neighbour trees

Neighbourhood tree species diversity might help to mitigate drought impacts due to complementarity effects, i.e. temporal or spatial resource partitioning that results in higher water availability in mixed than in pure communities, or facilitation, i.e. a positive effect on the functioning of cohabiting species (Ammer 2017; Anderegg et al. 2018, Grossiord 2019; Schnabel et al. 2019; Fichtner et al. 2020). According to our model, neighbourhood interspecific interactions played an important role for the survival of trees, either mitigating or enhancing drought stress depending on the surrounding species. This might explain controversial reports of species-mixing effects on drought stress. We used a modified competition index based on Hegyi (1974) to describe the effect of the relative height of each of the eight immediate neighbour trees on the probability of death of the focal tree (Fig. S1.1) while accounting for all possible inter- and intraspecific pairs of neighbour species constellations. We observed a negative neighbourhood effect on the risk of mortality for the majority of focal tree species (Fig. 5a). The number of neighbours of a particular species surrounding each focal tree varied depending on the species mixture in the plot and pre-drought background mortality. The presence of more and larger neighbours of a given species resulted in a reduced mortality risk in the majority of possible combinations. *A. platanoides* was the only species whose presence on average increased the mortality risk of its neighbours (Fig. 5a). Across pairs of tree species, beneficial interactions were much more common than detrimental ones (Fig. 5b). The correlation between the directed neighbourhood effects in a pair of neighbours was not credibly different from zero *(p_γ_*= −0.25, 95% HDI: −0.76 – 0.22), indicating that an advantage for one species in a pair did not always translate into a disadvantage for its neighbour. Instead, there was a tendency of pairs of neighbour species to be either one-sided or mutually beneficial, while not a single combination resulted in an increased mortality risk for both species (Fig. S2.4). In general, species that suffered more from drought increased the survival probability of their neighbours and vice versa. While *A. platanoides*, the species that most strongly increased the mortality of its neighbours, was among the species least affected by drought, four of the five species with positive effects on the survival of their neighbours suffered losses of over 50% (Fig. 1, Fig. 5a, Table 1). Besides an increased mortality of *P. pungens* in the neighbourhood of *P. abies,* the only species that reduced the survival of their neighbours were *A. platanoides* and *Q. rubra.* The presence of these two dominant overstory species increased the mortality especially for suppressed, subordinate species (Fig. 5). The increased survival of trees with neighbour species that suffered extreme losses may be a result of competitive release, especially in the case of *Larix* spp. and *Betula* spp., which did not compete for water after their early leaf shedding in 2018.

**Fig. 5.**
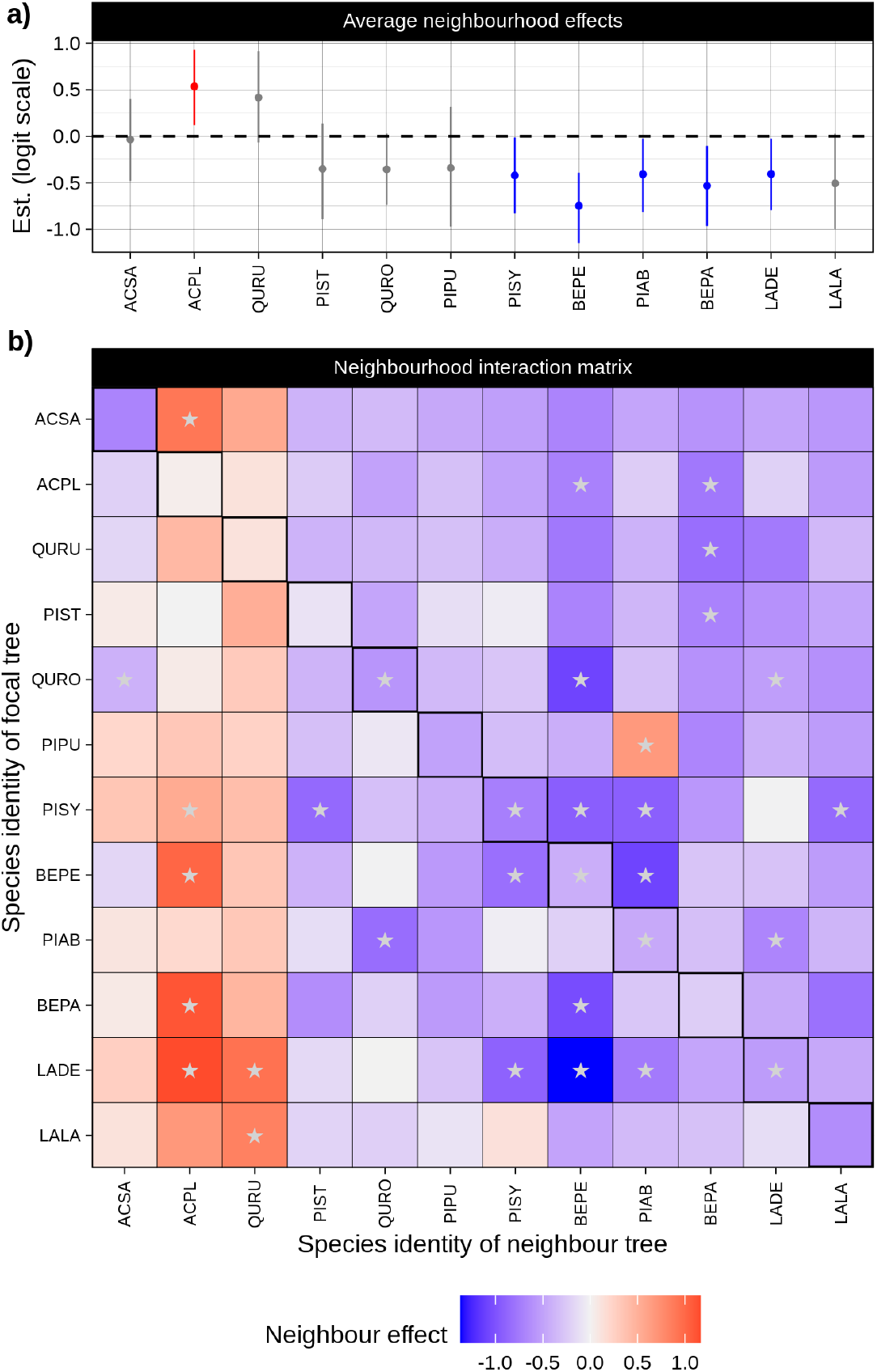
a) Estimated average neighbourhood effect on mortality on the logit scale (posterior mean ± 95% HDI). Blue: credibly reduced risk; red: credibly elevated risk; grey: no credible effect on mortality. b) Full matrix of the neighbourhood effects of a given species (columns) on focal trees of a given species (rows) on the logit scale. Colours indicate the strength of the effect; stars indicate effects that are credibly different from zero at the 95% level. The highlighted panels on the plot diagonal show intra-specific neighbourhood effects. Species are ordered by increasing mortality. For species acronyms, see Table 1.

An important implication of our model is that drought-related interactions between neighbouring trees are directed and species-specific, as previously observed for adult trees (Forrester et al. 2016). Studies of mitigation and competition tend to focus on effects of species richness (e.g. Fichtner et al. 2020), which integrate over all possible neighbourhood interactions. However, a net positive effect of species richness on survival only occurs if the interspecific neighbourhood interactions on average are more beneficial than intraspecific interactions. Whether or not there is a positive relationship with species richness therefore depends on the identity of the studied species rather than their mere number. Methods that preserve the information about the identity of interacting species allow for a more nuanced understanding of the underlying processes, and provide valuable additional information that e.g. permits identification of mutually beneficial species combinations. We hope that our approach to addressing neighbourhood effects contributes to a more thorough focus on the nature of directed neighbourhood interactions in plant drought responses, as in their sum they ultimately determine the resilience of a community against drought events.

### Nutrient and environmental effects

Conditional on the effect of the other predictor variables, none of the nutrient-enhanced treatments (N, P or NP addition) differed in mortality compared to the control treatment for any of the analysed species (Fig. 3, Fig. S2.2). Limitations posed by the design of the IDENT experiment preclude a clear interpretation of the nutrient effects, as fertilizer was only applied to the subset of European species, but not to the North-American ones, and several highly relevant species-level traits were only measured on the control treatment. So far, only a limited number of studies are available for the interaction between drought and plant nutrition and the role of nutrients in drought-induced mortality (Royo & Knight 2012; Wang et al. 2012; Sergent et al. 2014; Gessler et al. 2017). However, fertilization can reasonably be assumed to affect tree size, hydraulic efficiency and safety (Zhang et al. 2018), carbon dynamics (Li et al. 2018; Schönbeck et al. 2020) and susceptibility to bark beetles (Herms 2002), as well as other potential unmeasured predictors of mortality. The main rationale for including nutrient effects in our model was therefore not to accurately estimate their total effect on mortality, but to control for potential confounding effects (see Supplementary Material S1).

While no direct fertilization effects were observed, a considerable amount of the species-wise plot effects were credibly different from zero (Fig. S2.5), indicating the presence of small-scale environmental influences on tree survival. Though the precise origin of these differences is unknown, our results indicate they had a non-negligible contribution to the total variance in mortality risk, which may be associated with plot-specific differences, e.g. in soil texture, rooting depth or wind exposition.

### Conclusions

The extreme natural drought event in 2018 that killed over one third of the trees at the IDENT tree diversity experiment in Freiburg provided a unique opportunity to compare the mutually inclusive mechanisms underlying drought-induced mortality in one setting. While we support the view that severe tissue dehydration causing hydraulic failure marks the final stage in the process of tree death under drought, our results illustrate that hydraulic traits alone are not sufficient to predict individual mortality risk. Instead, the individual probability of reaching lethal desiccation thresholds depends on a series of mutually inclusive processes, notably starch depletion, known to be important for osmotic adjustment. Given the projected increase in drought exposure in most of the world’s forested biomes, improving the mechanistic understanding of processes involved in drought-induced tree mortality is essential. To predict future changes in the structure and composition of forests, it is crucial to adopt a holistic view on the drivers of tree death under drought. Specifically, there is a need for an improved understanding of the processes that link carbohydrate metabolism and water relations, and the ways mortality risk is modulated by pests, environmental conditions, and neighbourhood interactions among trees varying in size and species composition in a non-zero-sum list of effects.

## Methods

### Site description

The dataset for this study was collected in 2018/19 at the International Diversity Experiment Network with Trees (IDENT) in Freiburg, Germany (48°01’10”N, 7°49’37”E; 240 m a.s.l) (Tobner et al. 2014; Wein et al. 2016). IDENT belongs to the global network of tree diversity experiments (TreeDivNet, Paquette et al. 2018). Mean annual temperature was 11.6°C and mean annual precipitation 881.8 mm (long-term average from 1989-2019). Similar to the exceptional hot and dry conditions in large parts of Central Europe in 2018 (Hari et al. 2020; Schuldt et al. 2020), the IDENT study site was subjected to the most severe summer drought ever recorded in the region. The mean temperature from June to August in 2018 were 2.0°C above (20.9 vs. 18.9°C) and precipitation rates 62% below (118.2 vs. 314.2 mm) the long-term average from 1961-1990, respectively (DWD Climate Data Centre - CDC 2020a, b).

The study site comprises 415 plots of 13 m^2^, each containing 7 × 7 = 49 trees planted on a 0.45 × 0.45 m^2^ grid, resulting in a total of 20,335 trees. A buffer zone of 0.9 m separates neighbouring plots. The experiment manipulates tree diversity (richness,: monocultures and two, four, six species; functional diversity: mixtures of varying similarity in selected functional traits), geographic origin (European, North American), and species composition (different species mixtures per diversity level) in a four-times replicated block design using a species pool of six congeneric species pairs native to North America and Europe (Table 1). Mixtures comprising European species (with the exception of 6-species mixtures) were replicated for a tree species richness x fertilization sub-experiment with nitrogen (130 g elemental N per plot), phosphorous (65 g elemental P per plot) and N+P addition, respectively. All trees were planted in 2013 as 1-3 years old saplings on shallow sandy-loamy Cambisol (0.4 m) with a high gravel content bedrock (1.0 m).

### Tree mortality and height

We surveyed the survival and health status of all 20,335 planted trees at pre-drought (2017/18 inventory), directly after the drought in autumn 2018, and post-drought in early summer of 2019. Trees were classified as healthy, damaged (leaf browning or leaf loss), severely damaged (partial canopy dieback) or dead. Our analysis was based on the subset of the inner 25 trees of each plot that were alive during the pre-drought inventory, resulting in a total of 9,435 trees. These trees represent 90.9% of the initially planted individuals. Since boreholes of the bark beetle *Pityogenes chalcographus* were observed on several trees in early July 2018, we additionally recorded the beetle infestation for all conifers.

Pre-drought tree height (stem base to the apical branch) was measured during December 2017 and January 2018 for the 25 central trees of each plot as part of a yearly inventory. Additionally, a remote sensing campaign was carried out in July 2018 using a drone (Octo8XL, Mikrokopter GmbH, Saldenburg, Germany) equipped with an optical RGB camera with a 16 mm lens (Sony A5000, Stuttgart, Germany). The heights of the edge trees (for computation of neighbourhood effects) were then imputed based on these remote-sensing derived aggregate and rescaled by the average height of the corresponding species in each plot to account for species differences (see Supplementary Material S1 for details).

### Hydraulic traits

For measuring xylem embolism resistance, 100-150 cm long branch samples were collected from 12-19 live trees per species with the exception of the genus *Quercus* (3-5 trees × 4 plots × 10 species = 171 samples in total) in late summer 2019, immediately wrapped in moist towels and stored in black plastic bags to prevent dehydration during transport. Samples were re-cut under water directly before xylem vulnerability curves were constructed according to standard protocols for short-vesseled coniferous and diffuse-porous species using the flow-centrifuge technique (Cochard et al. 2005; Delzon et al. 2010; Schuldt et al. 2016). Vulnerability curves were fitted in R v4.0.0 (R Core Team 2020) with nonlinear least squares using a sigmoidal model based on the raw conductance measurements (Pammenter & Vander Willigen 1998; Ogle et al. 2009) (see details in Supplementary Material 1). Values for the long-vessel *Quercus* species were obtained from published data (Lobo et al. 2018).

Midday leaf water potential (*Ψ*_md_, MPa) was measured for four trees per species in all 48 monocultures at the peak of the drought (12-14 August) using a Scholander pressure chamber (1505D-EXP, PMS Instruments, Corvallis, USA).

Hydraulic safety margins (HSM) were computed as the difference between *Ψmd* and averages of the critical xylem pressure (*P*_crit_), which was assumed to be equivalent to the pressure at 88% loss of conductivity for angiosperms and 50% for conifers (Brodribb & Cochard 2009; Urli et al. 2013; Delzon & Cochard 2014). To account for the uncertainty in the estimated vulnerability curve parameters, it was propagated into the calculated HSM values (see Supplementary Material S1 for details).

### Non-structural carbohydrates

Leaf samples for the analysis of non-structural carbohydrate contents were taken during pre-drought (21 May to 18 June 2018) after leaf enfolding was completed in monocultures (48 plots), and post-drought (1–4 October 2018) on the full design, yielding ~1,000 sampled trees in total. Leaves were sampled from a random subset of the inner 5 × 5 trees representing the most common tree size whenever possible. Buffer trees were sampled if less than three trees per species were available for sampling in the core area. Each three to five leaves or 50–100 needles from a vital branch of the upper and middle part of the sun canopy were pooled, immediately stored at 4°C in the field and, within the same day, oven-dried at 60°C for 48 h to stop enzymatic activity (Landhäusser et al. 2018).

The low-molecular-weight sugars (glucose, fructose and sucrose) and starch were analyzed following the protocol of Wong (1990), modified according to Hoch et al. (2002) (see

Supplementary Material S1 for details). The change in relative sugar content (i.e. in the relative contribution of soluble sugars to the total non-structural carbohydrates) over the growing season (Δ_sugar_) was then calculated as the difference in the percentage of soluble sugars measured in May and in October 2018 in 48 unfertilized plots in the monocultures. The change in relative sugar content is expected to act as an indicator for the amount of starch-to-sugar conversion and depletion of starch reserves over the growing season (He et al. 2020).

### Neighbourhood effects

The influence of up to eight immediate neighbour trees on each of *i* in *I* central trees was expressed in form of a *I ×J* neighbourhood matrix ***N*** containing the sum of the heights *hj* of the trees of species *j* of *J* species divided by the height *hi* of the central tree, weighted by their relative distance *d_ij_* (i.e. with a weight of 1 for first and 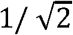 for second order neighbours; see Supplementary Material S1).

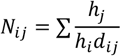

In this formulation, the computed value can be interpreted as a partial Hegyi competition index for a species (Hegyi, 1974), thus effectively decomposing the contribution of different competitor species to the total neighbourhood effect.

### Statistical modelling

In a hierarchical Bayesian modelling framework based on the Stan probabilistic programming language (Stan v. 2.21.0; Carpenter et al. 2017), we described the observed tree mortality status in 2019 as a function of the individual mortality risk.

We assumed that the observed state of a tree *(Y_Obs[ijk]_;* 0: tree alive, 1: tree dead) for each tree *i* belonging to species *j* in plot *k* for a total of *I* trees, *J* species and *K* plots is subject to observation errors with a species-specific probability of erroneously classifying a living tree as dead (e.g. because of leaf shedding). To account for this misclassification, we modelled the observed mortality state of individual trees as a one-inflated Bernoulli process with a tree-specific probability of dying *p_ijk_* and a species-specific misclassification probability *ϕ_j_*.

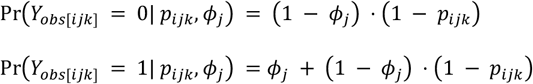

We further included observation models for the two species level traits in the focus of our main hypotheses, namely the hydraulic safety margin *HSM* and the shift in the fraction of soluble sugars of the total leaf non-structural carbohydrate content (Δ_sugar_) (see details in Supplementary Material S1).

We expressed the probability of dying of each tree as a logit-linear function of the true species-level Δ_sugar_ and *HSM* values, a term specifying species specific intercept and effects of tree height, bark beetle infestation and nutrient treatment *(I*×*L* predictor matrix ***X***), a term specifying neighbourhood effects *(I*×*J* neighbourhood matrix ***N***, elevated to a power of *c*), and species-specific plot effects ***δ_jk_***.

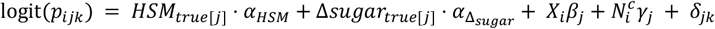

The *J*×*L* species-specific regression coefficients ***β*** were described by a multivariate normal distribution with covariance matrix ***Σ*_β_**:

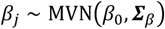

To account for a possible correlation between the directed neighbourhood effects within a pair of species, the *J*×*J* neighbourhood effects matrix ***γ*** for focal species *m* and neighbour species *n* was parameterized as follows:

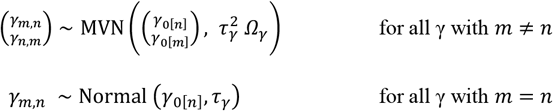

where *Ω_γ_* is a 2 × 2 correlation matrix allowing for a correlation of the directed neighbourhood effects in pairs of non-identical species (the off-diagonal elements of ***γ***).

The design effects δ for each plot and species were modelled by a normal distribution centred around zero with standard deviation *τ_δ_*, i.e. as a varying intercept for each combination of plot and species:

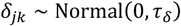

Modelling was performed via R using rstan v. 2.19.3 (Stan Development Team, 2020). The model reached a *Ȓ* statistic of under 1.002 and effective sample sizes above 1,500 for all estimated parameters, indicating full model convergence (Table S2.1). The estimated species-specific misclassification rates *ϕ* were low (on average 0.9%; Supplementary Material S1), though higher rates of up to 4.2% of trees erroneously classified as dead were found for some species (Table S2.1, Fig. S2.1), resulting in a visible mismatch between observed and predicted mortality for these species in Fig. 1. Details about model structure, prior specification, model implementation and model fitting are provided in Supplementary Material S1. The model code and raw data are available as a repository on Github (Supplementary Material S3).

## Supporting information

Supplementary Results

## Acknowledgements

We thank C. Gernert, Y. Heppenstiel, B. Kukatsch, S. Ouyang and L. Schönbeck for their support in the laboratories and S. Bilodeau-Gauthier, A. Böminghaus, G. Csapek, A.-L. Hillebrecht, A. Klingler, D. Saito, A. Schäfer, D. Sprenger, M. Romer, M. Ulmerich, J. Maron and M. Witt for their contribution to data collection at IDENT Freiburg. Funding by the German Research Foundation (DFG 384026712 to LR; DFG 316733524 to CN) and the University of Freiburg (Innovationsfonds Forschung to MSL and JB) is gratefully acknowledged. The research presented here contributes to the International Tree Mortality Network (https://www.tree-mortality.net/), an initiative of the IUFRO Task Force on Tree Mortality (https://www.iufro.org/science/task-forces/tree-mortality-patterns/), and to the Global Network of Tree Diversity Experiments (TreeDivNet; https://treedivnet.ugent.be/), belonging to the IDENT network within TreeDivNet.

## Author contributions

A.P. and C.M. developed the original IDENT design. M.S.-L. and. J.B. expanded and implemented the design for the Freiburg trial and maintained it together with C.N. and T.G.. L.R., B.S. and A.G. developed the presented study. C.N. was responsible for pre-drought inventories, P.H. performed the mortality inventory and field sampling with support from L.R., and K.K. provided drone derived tree heights. P.H. performed all physiological measurements with support from M.S.. R.M.L. developed the model and performed the statistical analyses, and P.H., R.M.L. and B.S. wrote the first version of the manuscript, which was intensively discussed and revised by all authors.

## Supplementary material

S1: Supplementary methods

S2: Supplementary results

S3: Digital supplement: A repository containing the code for data processing and fitting the model described in S1 and the full dataset is available upon request and will be made public after publication of the article under the following address: https://github.com/r-link/mutually_inclusive_mechanisms

