## Supplementary Methods for "Mutually inclusive mechanisms of drought-induced tree mortality"

### Supplementary material 1: Supplementary methods

Supplementary material for Hajek et al. 2020

#### Xylem vulnerability curves

Xylem vulnerability curves were described with the conductance-based reparameterization of the logistic model of Pammenter and Vander Willigen (1998) proposed by Ogle et al. (2009) and fitted with with nonlinear least squares using base R function `nls()`:

$$K_i \sim \text{Normal}\left(k_{sat}\left(1 - \frac{1}{1 + \exp\left(s/25\left(\Psi_i - \Psi_{50}\right)\right)}\right),\ \sigma\right) \tag{1}$$

,

where $K_i$ is the raw hydraulic conductivity for measurement $i$ as measured with the Cavitron, $\Psi_i$ is the corresponding xylem pressure, $k_{sat}$ is the (unknown) maximum conductivity at full saturation of the sample, $\Psi_{50}$ is the xylem pressure at 50% loss of conductivity, $s$ is the (negative) slope of the curve at the $\Psi_{50}$, and $\sigma$ is the residual standard deviation. The xylem pressure at 12% and 88% loss of conductivity ($\Psi_{12}$ and $\Psi_{88}$, respectively) were then calculated by expressing the model equation in terms of PLC, fixing PLC to 12% or 88% and solving for $\Psi$:

$$\Psi_{12} = \frac{25\ \ln\left(\frac{100}{12} - 1\right)}{s} + \Psi_{50} \qquad \Psi_{88} = \frac{25\ \ln\left(\frac{100}{88} - 1\right)}{s} + \Psi_{50}$$

Approximate standard errors and confidence intervals for these quantities were obtained by sampling from the models' covariance matrices (cf. "population prediction intervals" in Lande et al. 2003; Bolker 2008).

#### Accounting for the uncertainty in hydraulic traits

Due to technical limitations, plant hydraulic measurements could only be performed on a subset of trees. Moreover, vulnerability curves and leaf water potentials were not measured on the same tree individuals, which impeded calculating hydraulic safety margins for individual trees. For that reason, we decided to treat the hydraulic safety margins as a species-level trait. We accounted for the uncertainty in the estimated hydraulic traits by propagating it into the species-level aggregates of these variables.

To achieve this, weighted species averages of $\Psi_{crit}$ ($\Psi_{50}$ or $\Psi_{88}$, approximate lethality threshold for gymnosperm/angiosperm trees, respectively; cf. Brodribb and Cochard 2009; Urli et al. 2013; Blackman et al. 2016) were calculated using inverse variance weights based on the standard errors of the estimates.

$$\overline{\Psi_{crit}} = \sum_{i=1}^{n} \frac{\frac{1}{SE_i^2}\Psi_{crit[i]}}{\sum_{j=1}^{n}\frac{1}{SE_j^2}} \tag{2}$$

This estimate is equivalent to the mean effect size in a fixed-effects meta-analytic model. The standard error of $\overline{\Psi_{crit}}$ can be calculated by the following equation (Rosenberg et al. 2013):

$$SE\left(\overline{\Psi_{crit}}\right) = \sqrt{\frac{1}{\sum_{i=1}^{n}\frac{1}{SE_i^2}}} \tag{3}$$

Hydraulic safety margins were then calculated as the difference between the species averages of the mid-day water potentials in August 2018 ($\overline{Psi_{md}}$) and the species estimate for$\overline{\Psi_{crit}}$:

$$HSM = \overline{\Psi_{md}} - \overline{\Psi_{crit}} \tag{4}$$

The standard error of these estimates is given by the equation for the variance of a difference of random variables:

$$SE(HSM) = \sqrt{SE\left(\overline{\Psi_{crit}}\right)^2 + SE\left(\overline{\Psi_{md}}\right)^2} \tag{5}$$

#### Chemical analysis of non-structural carbohydrates

The contents of low-molecular-weight sugars (glucose, fructose and sucrose) as well as the starch content were analyzed with a modification of the protocol of Wong (1990) according to Hoch, Popp, and Korner (2002). 10 mg of the grinded leaf material was boiled in 2 ml of distilled water for 30 min. After centrifugation, an aliquot of 200 µl was incubated with invertase and isomerase from baker's yeast (I4504 and P5381 Sigma-Aldrich) to degrade sucrose and convert fructose into glucose. The total amount of glucose (sugars) was determined photometrically at 340 nm in a 96-well microplate photometer (HR 7000; Hamilton, Reno, NE, USA) after enzymatic conversion to gluconate-6-phosphate (GHK assay reagent, I4504 and G3293, Sigma-Aldrich). The total amount of NSC was measured by taking 500 µl of the extract (including sugars and starch) incubated with a fungal amyloglucosidase from Aspergillus niger (10115 Sigma-Aldrich) for 15h at 49°C to digest starch into glucose. Total glucose was determined photometrically as described above. The concentration of starch was calculated as NSC minus free sugars. Pure starch and glucose, fructose and sucrose solutions were used as standards, and standard plant powder (Orchard leaves; Leco, St Joseph, MI, USA) was included to control the reproducibility of the extraction. NSC concentrations are expressed on a percentage dry matter basis.

#### Construction of the neighbourhood matrix

The influence of the eigh neighbour trees of first and second order (cf. Fig. S1.1) on each of $i$ in $I$ focal trees was expressed in form of a $I \times J$ neighbourhood matrix containing the sum of the heights $h_j$ of the trees of species $j$ of $J$ species divided by the height $h_i$ of the central tree, weighted by their relative distance $d_{ij}$ (i.e. with a weight of 1 for first and $1/\sqrt{2}$ for second order neighbours).

$$N_{ij} = \sum \frac{h_j}{h_i d_{ij}} \tag{6}$$

The value of $N_{ij}$ can be interpreted as a partial Hegyi competition index (Hegyi 1974) describing the competitive pressure of immediate neighbour trees belonging to a particular species on the focal tree. Accordingly, our approach decomposes the neighbourhood effect into the contributions of different contributor species, and therefore allows for different types of neighbourhood interactions depending on the corresponding species pairs.

#### Drone-based height estimates for the buffer trees

To compute the neighbourhood effects for the outermost line of the 25 central trees based on Equation (6), it is necessary to have height estimates of the buffer trees. As direct tree height measurements were only available for the 25 central trees of each plot, for this purpose the heights of the edge tree were imputed based on the remote-sensing derived aggregate heights for the plot margins and corners estimated from drone-based aerial imaging.

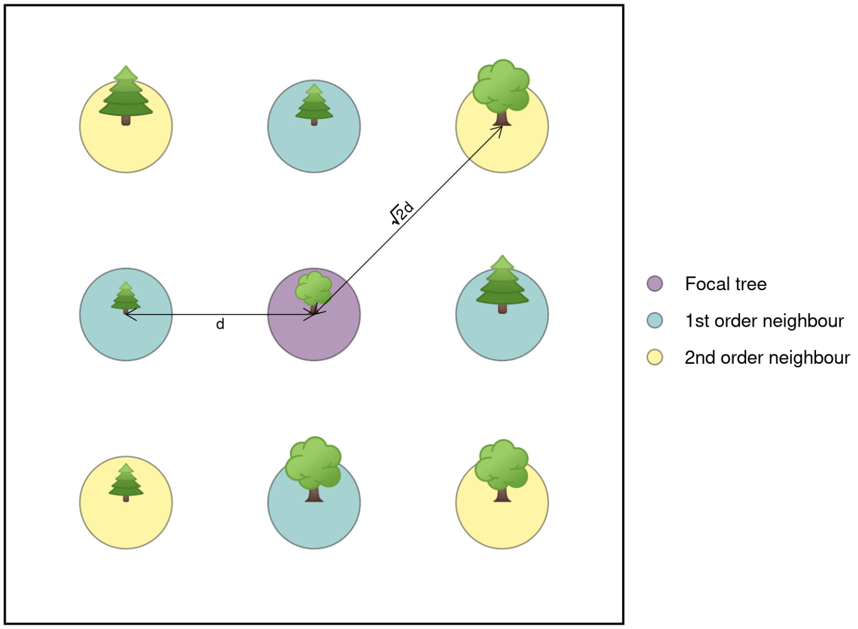


**Figure S1.1** Schematic drawing of a focal tree and its eight immediate neighbours.

A sitewide drone flight campaign was carried out on July 20, 2018 using a Octo8XL drone (Mikrokopter GmbH, Saldenburg, Germany) fitted with a Sony A5000 RGB camera (Sony A5000, Stuttgart, Germany) with a 16 mm lens (24mm @ 35 mm equivalent) with remote trigger operation. 20 ground control markers were systematically laid out across the IDENT experiment and their precise geographic position was determined using a Leica GNSS GS16 RTK unit. The drone was then flown using autopilot missions at 45 meters above ground level on an image capture interval that produced approximately 90% forward photo overlap and 80% sidelap. 370 photos were then processed into structure from motion models using high quality settings in Agisoft Metashape (Agisoft LLC, St. Petersburg, Russia), and a 2.62cm/pix digital elevation model was rendered. Using the corresponding 1.31cm/pix RGB orthomosaic, specific polygons representing plot core trees were used to create buffers of edge trees in both the four cardinal directions and four plot corners. From these buffers, mean sea level was extracted from the digital elevation model and subtracted from plot core elevation to derive relative buffer and core height differences.

As the heights of individual trees could not reliably be resolved based on the drone-based height estimates, they did not contain information about the (often large) differences in average size between the different species in a plot. To avoid losing the information about species-based size differences, the drone-based height estimates in multi-species plots were rescaled by the differences of the average height of the corresponding species in each plot from the overall mean height in the plot.

#### Model structure

##### Directed acyclic graphs

In order to identify potential confounders whose exclusion may result in bias the estimated effects of the central predictor variables in our model, as an initial step of the modelling process we built a set of directed acyclic graphs (DAGs) of the assumed causal relationships between these variables. We assumed HSM, changes in NSC pools, bark beetle infestation, tree size and neighbourhood effects to all have a direct effect on mortality. In addition, we supposed that the size of a tree will both affect its ability to build up NSC pools and its susceptibility to bark beetles, and moreover is by definition associated with the (size-dependent) neighbourhood effects. We further assumed that HSMs and changes in NSC pools are associated, and that species with smaller HSM are at a higher risk of beetle infestation (Fig. S1.2).

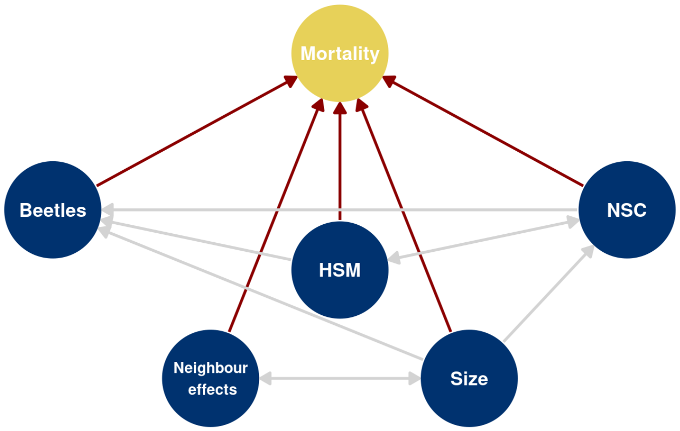


**Figure S1.2** Directed acyclic graph with the main predictors of mortality considered in this work and their expected interellationships.

A complication for our model is that bark beetle infestation was only recorded once during the main drought event in 2018. As we could not rule out a further spread of the infestation starting from the trees affected at the time of measurement, we assumed that an effect of the presence of bark beetles in the immediate neighbourhood of the focal tree on its survival was possible. In addition, as a part of the IDENT experiment, a number of plots was subjected to a fertilization treatment, and plots could be assumed to differ slightly in other environmental variables. Both nutrient treatment and random plot differences in environmental conditions can be expected to have an effect on most of the central predictor variables, and are most likely associated with other, unmeasured plant properties associated with the probability of survival (Fig. S1.3).

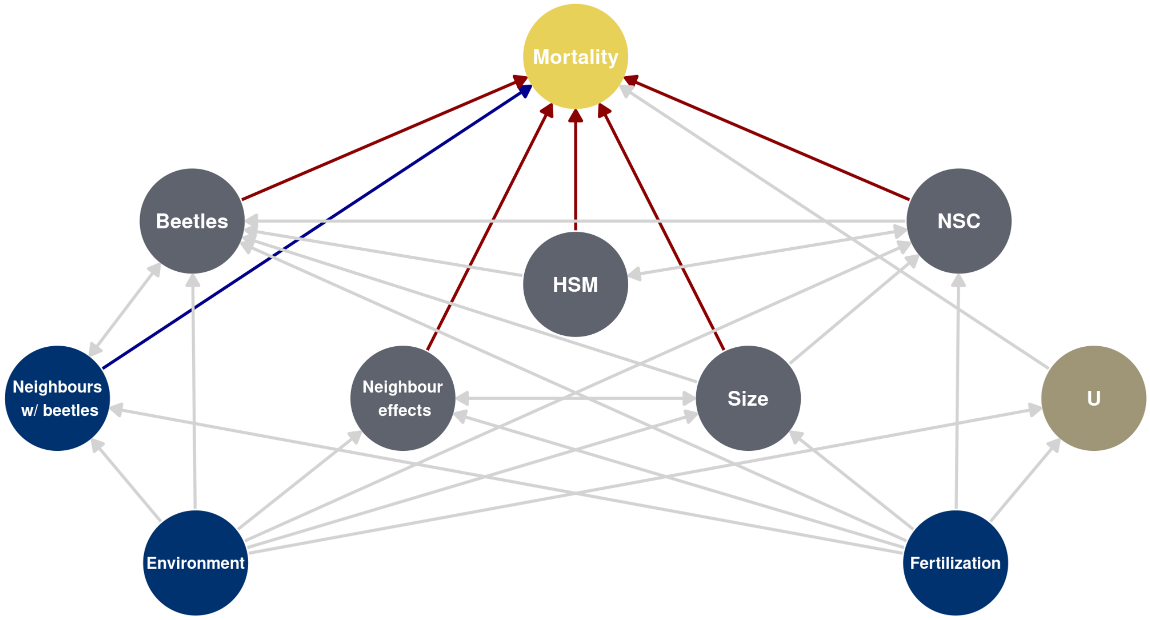


**Figure S1.3** Extended directed acyclic graph including the effect of neighbour-trees infected by bark beetles, and taking into account potential confounders, i.e. small-scale environmental differences, nutrient treatment and unmeasured variables (U).

These unmeasured variables open a potential backdoor path via the environmental variables and nutrient

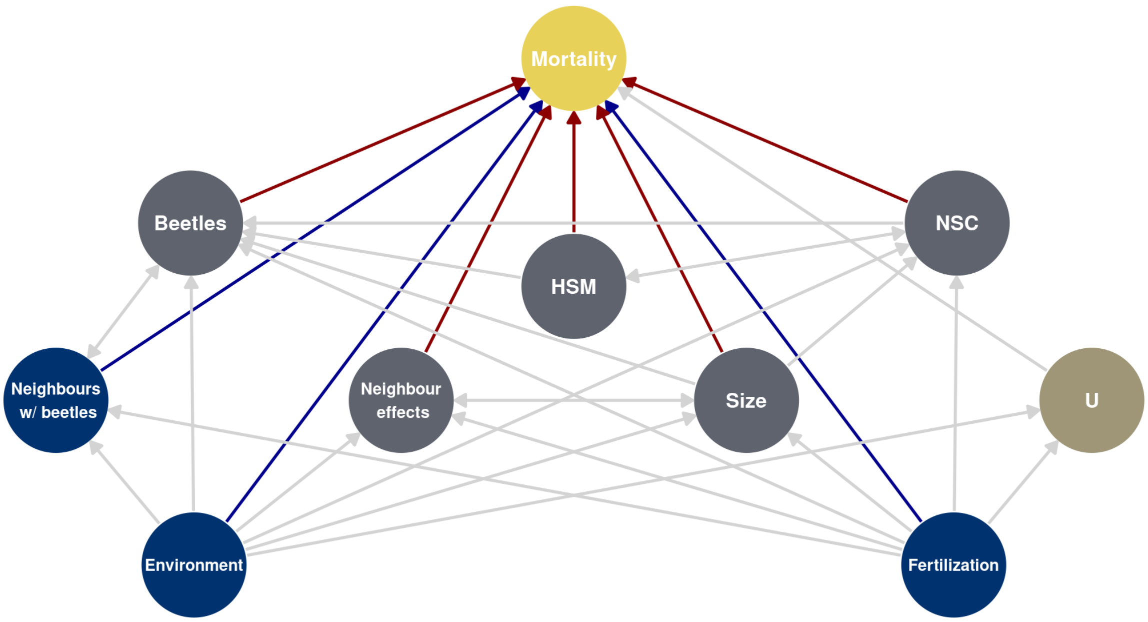


**Figure S1.4** Same DAG containing covariates necessary to close backdoor paths of environmental variables, nutrient treatment and unmeasured variables on mortality.

treatment, potentially resulting in omitted variable bias (cf. Shrier and Platt 2008; VanderWeele, Hernán, and Robins 2008). To close this backdoor path, it was necessary to include predictors for the fertilization treatment and for environmental differences between plots (cf. Fig. S1.4), even though we did not expect strong effects of fertilization on mortality.

We are aware that due to the observational nature of our dataset with all its shortcomings, a strict causal interpretation of our model is not warranted. However, we believe that adopting a causal logic and using of graph theoretical tools like DAGs may help to identify potential sources of bias and allows for more informed model building.

#### Observation model

In our model, we described the observed tree mortality status in 2019 $Y_{obs[ijk]}$ (0: tree survived, 1: tree dead) for each tree $i$ belonging to species $j$ in plot $k$ for a total of $I$ trees, $J$ species and $K$ plots as a function of an individual mortality risk. We assumed that the classification of a tree as dead or alive is subject to observation errors with a probability of erroneously classifying a living tree as dead (e.g. because of leaf shedding) with a species-specific misclassification probability $\phi_j$.

For this reason, we modeled the observed mortality state of individual trees as a 'one-inflated' Bernoulli process with a tree-specific probability of dying $p_{ijk}$.

$$\Pr\left(Y_{obs[ijk]} = 0 \mid p_{ijk},\, \phi_j\right) = (1-\phi_j)\cdot(1-p_{ijk}) \tag{7}$$

$$\Pr\left(Y_{obs[ijk]} = 1 \mid p_{ijk},\, \phi_j\right) = \phi_j + (1-\phi_j)\cdot p_{ijk} \tag{8}$$

We further included observation models for the two species level traits in the focus of our main hypotheses, namely the hydraulic safety margin $HSM$ and the shift in the fraction of soluble sugars of the total leaf nonstructural carbohydrate content (sugar fraction in October minus sugar fraction in May; in the following abbreviated as $\Delta sugar$).

For the hydraulic safety margin, we assumed that the observed values $HSM_{obs[j]}$ (scaled to a common mean of zero and a standard deviation of one) were distributed normally around the true population averages $HSM_{true[j]}$ for that species with a fixed standard deviation given by the standard error $SE_{HSM[j]}$ for the estimate of the species in question:

$$HSM_{obs[j]} \sim \text{Normal}\left(HSM_{true[j]},\, SE_{HSM[j]}\right) \tag{9}$$

For the change in relative sugar content, we assumed that the (scaled and centered) observed values $\Delta sugar_{obs[jt]}$ for each species were normally distributed around the true species-level population averages $\Delta sugar_{true[j]}$ with a common standard deviation $\sigma_{NSC}$.

$$\Delta sugar_{obs[jt]} \sim \text{Normal}\left(\Delta sugar_{true[j]},\, \sigma_{\Delta sugar}\right) \tag{10}$$

The misclassification parameters $\phi$ were assigned moderately informative Beta($a = 0.1, b = 1$) priors and constrained to values below 0.5 to assure the identifiability of the model.

$$\phi_j \sim \text{Beta}(0.1, 1) \tag{11}$$

The latent true values $HSM_{true}$ and $\Delta sugar_{true}$ were both assigned a standard normal prior. The standard deviation of the individual $\Delta sugar$ differences $\sigma_{\Delta sugar}$ was modeled with an informative Normal$\,(0,\, 0.2)$ prior (equivalent to the prior assumption that the between-species standard deviation in $\Delta sugar$ is 5 times larger than the within-species standard deviation).

$$HSM_{true[j]} \sim \text{Normal}\,(0,\, 1) \tag{12}$$

$$\Delta sugar_{true[j]} \sim \text{Normal}\,(0,\, 1) \tag{13}$$

$$\sigma_{\Delta sugar} \sim \text{Normal}\,(0,\, 0.2) \tag{14}$$

##### Process model

We expressed the probability of dying $p_{ijk}$ of each tree as a logit-linear function of the true species-level $\Delta sugar$ and $HSM$ values, a term specifying species specific size, trait and treatment effects, ($I \times L$ predictor matrix $\mathbf{X}$), a term specifying neighbourhood effects ($I \times J$ neighbourhood matrix $\mathbf{N}$), and a term $\boldsymbol{\delta}_{jk}$ specifying design effects.

$$\text{logit}\,(p_{ijk}) = HSM_{true[j]} \cdot \alpha_{HSM} + \Delta sugar_{true[j]} \cdot \alpha_{\Delta sugar} + \mathbf{X}_i \boldsymbol{\beta}_j + \mathbf{N}_i^c \boldsymbol{\gamma}_j + \boldsymbol{\delta}_{jk} \tag{15}$$

**Size, herbivore and nutrient effects** The predictor matrix $X$ contained 1) a constant intercept term, 2) a linear term for log-transformed, scaled and centered tree size, indicator variables for 3) the presence of bark beetles in tree $i$ and 4) the presence of bark beetles in any of its eight immediate neighbour trees as detected in the summer of 2018, as well as indicators for the nutrient treatment, i.e. addition of 5) nitrogen, 6) phosphorous, or 7) both. As none but two species (*Larix decidua* and *Picea abies*) showed substantial levels of beetle infestation which led to estimation problems if performed on the species level, for this variable only the main effect $\beta_0$ was included in the model (i.e. the species-specific variation of the parameters for these variables was constrained to zero).

The species specific size, bark beetle and nutrient effects $\boldsymbol{\beta}$ were assumed to follow a multivariate normal distribution with overall mean vector $\boldsymbol{\beta}_0$ and covariance matrix $\boldsymbol{\Sigma}_\beta$.

$$\boldsymbol{\beta}_j \sim \text{MVN}\,(\beta_0,\, \boldsymbol{\Sigma}_\beta) \tag{16}$$

For the ease of interpretation and choice of priors, the covariance matrix of the species effects was further decomposed into a vector of standard deviations $\boldsymbol{\tau}_\beta$ and a correlation matrix $\boldsymbol{\Omega}_\beta$:

$$\boldsymbol{\Sigma}_\beta = \text{diag}\left(\boldsymbol{\tau}_\beta\right)\, \boldsymbol{\Omega}_\beta\, \text{diag}\left(\boldsymbol{\tau}_\beta\right) \tag{17}$$

**Neighbourhood effects** As a pragmatic way to reduce the effect of single trees with very extreme values in their neighbourhood matrix $\mathbf{N}$ (mostly very small trees with large neighbours, leading to extreme values in Eqn. (6) due to the small values in the denominator), the neighbourhood matrix was transformed by elevating it to a power of $c$ which was estimated from the data.

The effect of the relative size of the species $m$ on the species $n$ ($\gamma_{m,n}$) and vice versa ($\gamma_{n,m}$) were expressed in form of a $J \times J$ matrix $\boldsymbol{\Gamma}$. The neighbourhood effects for all possible species pairings were described as species-specific deviations from the overall average neighbourhood effect resulting from a neighbour of species $m$ $\gamma_{0[n]}$. The off-diagonal elements of $\gamma$ were drawn from a multivariate normal distribution with a mean vector given by $\gamma_0$ and a covariance matrix $\Sigma_\gamma$ to allow for a correlation between the directed interactions in the same species pair:

$$\begin{pmatrix} \gamma_{m,n} \\ \gamma_{n,m} \end{pmatrix} \sim \text{MVN}\left(\begin{pmatrix} \gamma_{0[n]} \\ \gamma_{0[m]} \end{pmatrix}, \boldsymbol{\Sigma}_\gamma\right) \quad \text{for all } \gamma \text{ with } m \neq n \tag{18}$$

The covariance matrix was again decomposed into a correlation matrix $\boldsymbol{\Omega}_\gamma$ and a standard deviation $\tau_\gamma$, though in this case assuming a constant standard deviation for both groups.

$$\boldsymbol{\Sigma}_\gamma = \tau_\gamma^2\, \boldsymbol{\Omega}_\gamma \tag{19}$$

For the diagonal elements of $\boldsymbol{\Gamma}$, the neighbourhood effects were expressed with a univariate normal distribution with the same standard deviation $\tau_\gamma$.

$$\gamma_{m,n} \sim \text{Normal}\left(\gamma_{0[n]}, \tau_\gamma\right) \quad \text{for all } \gamma \text{ with } m = n \tag{20}$$

**Design effects** The design effects were described as a varying effects term for each different combination of species and plot and were assumed to be normally distributed around 0 with a standard deviation $\tau_{plsp}$. As there were only three blocks which are all situated in close spatial proximity, it was assumed that potential block-specific differences were sufficiently subsumed in the species-specific plot effects:

$$\delta_{jk} \sim \text{Normal}\left(0,\, \tau_\delta\right) \tag{21}$$

##### Prior specifications

The main effects $\alpha_{HSM}$, $\alpha_{\Delta sugar}$, $\beta_{0[2\cdots L]}$ and $\gamma_0$ were assigned weakly regularizing normal priors with a mean of 0 and a standard deviation of 1.5, which are largely flat on the (inverse-logit transformed) response scale (cf. page 328 in McElreath 2020).

$$\alpha_{HSM} \sim \text{Normal}\left(\mu = 0,\, \sigma = 1.5\right) \tag{22}$$

$$\alpha_{\Delta sugar} \sim \text{Normal}\left(\mu = 0,\, \sigma = 1.5\right) \tag{23}$$

$$\boldsymbol{\beta} \sim \text{Normal}\left(\mu = 0,\, \sigma = 1.5\right) \tag{24}$$

$$\boldsymbol{\gamma}_0 \sim \text{Normal}\left(\mu = 0,\, \sigma = 1.5\right) \tag{25}$$

The coefficient $c$ controlling the skew of the neighbourhood effects was assigned a log-normal prior with a mean of 0 and a standard deviation of 1 on the log scale.

$$\log(c) \sim \text{Normal}\left(\mu = 0,\ \sigma = 1\right) \tag{26}$$

The standard deviations of the varying species and neighbourhood effects as well as the species-specific plot effects were described with half t-priors with a mean of 0, a standard deviation of 1.5 and 3 degrees of freedom, thus placing the bulk of probability mass in the low range while not ruling out occasional larger values:

$$\boldsymbol{\tau}_\beta \sim \text{half-t}\left(\mu = 0,\ \sigma = 1.5,\ \nu = 3\right) \tag{27}$$

$$\tau_\gamma \sim \text{half-t}\left(\mu = 0,\ \sigma = 1.5,\ \nu = 3\right) \tag{28}$$

$$\tau_\delta \sim \text{half-t}\left(\mu = 0,\ \sigma = 1.5,\ \nu = 3\right) \tag{29}$$

The correlation matrices of the varying effects $\boldsymbol{\Omega}_\beta$ and $\boldsymbol{\Omega}_\gamma$ were modeled with the so-called LKJ prior (i.e. the "extended onion method" of Lewandowski, Kurowicka, and Joe 2009) with a scale of 2.

$$\boldsymbol{\Omega}_\beta \sim \text{LKJcorr}(2) \tag{30}$$

$$\boldsymbol{\Omega}_\gamma \sim \text{LKJcorr}(2) \tag{31}$$

The LKJcorr(2) prior is a multivariate generalization of the boundary avoiding Beta$(2, 2)$ prior for correlation parameters recommended on page 317 of Gelman et al. (2014), and hence can be assumed to rule out extreme values of correlations of -1 or 1 while not contradicting any other possible values.

#### Model implementation details

The model was implemented in the Stan probabilistic programming language (Stan v. 2.21.0; Carpenter et al. 2017) using R package `rstan` v. 2.19.3 (Stan Development Team 2020) accessed through R v. 4.0.0 (R Core Team 2020).

Stan performs Markov Chain Monte Carlo (MCMC) sampling using Hamiltonian Monte Carlo (HMC) based on the no-U-Turn sampler (Hoffman and Gelman 2014). A number of reparameterizations and alternative expressions for certain model components can be exploited to increase the computational speed and accelerate convergence in HMC models. In our model, we took advantage of the so-called non-centered parameterization (Betancourt and Girolami 2015) to express $\boldsymbol{\delta}$, i.e. we decomposed the varying plot effects into a sample from a standard normal deviate, which was then post-multiplied with the corresponding scale in a later step. Analogously, the species-specific varying effects on $\boldsymbol{\beta}$ and $\boldsymbol{\gamma}$ were expressed in the multivariate generalization of the non-centered parameterization by drawing samples from independent standard normal deviates and then multiplying them with the corresponding covariance matrix. In addition, for computational reasons, the LKJ prior was put on the Cholesky factor of the corresponding correlation matrices instead of the correlation matrix itself.

The code for the model was developed based on modified code from a simplified model which could be expressed with the `brms` package (Bürkner 2017, 2018), which we also exploited to pre-process the data for model fitting (cf. code in the Github repository in Supplementary Material 3).

Starting values were fixed for the latent true values for *NSC* and *HSM* (started from their sample averages), for the between-species standard deviation for the *NSC* measurement model (started from $\sigma_{NSC} = 0.1$), the misclassification parameters (started from $\phi_j = 0.001$), and the scaling parameter for the neighbourhood matrix (started from $c = 0.8$). All other model parameters were started from random starting values as per the Stan default settings. The sampling was run for 5000 iterations (2500 of which were discarded as a burn-in interval) on 4 chains using a maximum tree depth of 15 and a target acceptance rate of `adapt_delta = 0.95`.

#### Model checking

Model convergence was assessed based on the bulk and tail effective sample size, the rank-based improved $\hat{R}$ convergence statistic by Vehtari et al. (2020), as well as by visual inspection of trace plots of the most relevant model parameters.

To assess the model's sensitivity to extremely influential observations and identify model misspecification, we performed approximate leave-one-out cross-validation based on Pareto-smoothed importance sampling (Vehtari, Gelman, and Gabry 2017; Yao et al. 2017) using R package `loo` (Vehtari et al. 2019). We made use of the shape parameter $k$ of the Pareto distribution to identify potentially influential observations. While initial model specifications had a number of observations with $k \geq 0.7$ (indicating that the posterior distribution cannot be effectively explored for these observations), the problem could be strongly alleviated after the introduction of the scaling parameter $c$ for the neighbourhood matrix, which downweighted the effect of extreme neighbourhood effects resulting for occasional neighbour pairs with very large size differences. In consequence, the final model specification with $c$ resulted in zero observations with $k \geq 0.7$ and only 21 observations with $0.7 > k \geq 0.5$, indicating that the introduction of the scaling parameter resolved the misspecification issues.

#### Explained variance

The variance in the observed tree mortality that was explained by the model was approximated with the Bayesian R-squared proposed by Gelman et al. (2018) as the variance of the predicted values for each simulation draw divided by the sum of the variance of the predicted values and the variance of the residuals in the respective draws:

$$R^2_{Bayes} = \frac{\mathrm{Var}(\hat{Y}_i)}{\mathrm{Var}(\hat{Y}_i) + \mathrm{Var}(Y_i - \hat{Y}_i)} = \frac{\mathrm{Var}\left(\Pr\left(Y_{obs[ijk]} = 1 \mid p_{ijk}, \phi_j\right)\right)}{\mathrm{Var}\left(\Pr\left(Y_{obs[ijk]} = 1 \mid p_{ijk}, \phi_j\right)\right) + \mathrm{Var}\left(Y_{obs[ijk]} - \Pr\left(Y_{obs[ijk]} = 1 \mid p_{ijk}, \phi_j\right)\right)} \tag{32}$$

The reported pseudo-R-squared value for our model is the posterior mean of this quantity.

Bolker, Benjamin M. 2008. *Ecological Models and Data in R*. Princeton and Oxford: Princeton University Press.

Brodribb, Tim J., and Hervé Cochard. 2009. "Hydraulic Failure Defines the Recovery and Point of Death in Water-Stressed Conifers." *Plant Physiology* 149 (1): 575–84. https://doi.org/10.1104/pp.108.129783.

Bürkner, Paul-Christian. 2017. "brms: An R Package for Bayesian Multilevel Models Using Stan." *Journal of Statistical Software* 80 (1): 1–28. https://doi.org/10.18637/jss.v080.i01.

———. 2018. "Advanced Bayesian Multilevel Modeling with the R Package brms." *The R Journal* 10 (1): 395–411. https://doi.org/10.32614/RJ-2018-017.

Carpenter, Bob, Andrew Gelman, Matthew D. Hoffman, Daniel Lee, Ben Goodrich, Michael Betancourt, Marcus Brubaker, Jiqiang Guo, Peter Li, and Allen Riddell. 2017. "Stan : A Probabilistic Programming Language." *Journal of Statistical Software* 76 (1). https://doi.org/10.18637/jss.v076.i01.

Gelman, Andrew, John B Carlin, Hal S Stern, David B Dunson, Aki Vehtari, and Donald B Rubin. 2014. *Bayesian Data Analysis*. 3rd ed. Boca Raton, FL: Chapman & Hall.

Gelman, Andrew, Ben Goodrich, Jonah Gabry, and Aki Vehtari. 2018. "R-Squared for Bayesian Regression Models." *The American Statistician*, December, 1–6. https://doi.org/10.1080/00031305.2018.1549100.

Hegyi, Frank. 1974. "A Simulation Model for Managing Jack-Pine Stands." In *Growth Models for Tree and Stand Simulation*, edited by Joran Fries, 30:74–90. Stockholm, Sweden: Royal College of Forestry.

Hoch, Gunter, Marianne Popp, and Christian Korner. 2002. "Altitudinal Increase of Mobile Carbon Pools in Pinus Cembra Suggests Sink Limitation of Growth at the Swiss Treeline." *Oikos* 98 (3): 361–74. https://doi.org/10.1034/j.1600-0706.2002.980301.x.

Hoffman, Matthew D, and Andrew Gelman. 2014. "The No-U-Turn Sampler: Adaptively Setting Path Lengths in Hamiltonian Monte Carlo." *Journal of Machine Learning Research* 15 (1): 1593–1623.

Lande, Russell, Steinar Engen, Bernt-Erik Saether, and others. 2003. *Stochastic Population Dynamics in Ecology and Conservation*. Oxford, England: Oxford University Press.

Lewandowski, Daniel, Dorota Kurowicka, and Harry Joe. 2009. "Generating Random Correlation Matrices Based on Vines and Extended Onion Method." *Journal of Multivariate Analysis* 100 (9): 1989–2001.

McElreath, Richard. 2020. *Statistical Rethinking: A Bayesian Course with Examples in R and Stan*. 2nd ed. CRC Press.

Ogle, Kiona, Jarrett J. Barber, Cynthia Willson, and Brenda Thompson. 2009. "Hierarchical Statistical Modeling of Xylem Vulnerability to Cavitation." *New Phytologist* 182 (2): 541–54. https://doi.org/10.1111/j.1469-8137.2008.02760.x.

Pammenter, N. W., and C. Vander Willigen. 1998. "A Mathematical and Statistical Analysis of the Curves Illustrating Vulnerability of Xylem to Cavitation." *Tree Physiology* 18 (8-9): 589–93. https://doi.org/10.1093/treephys/18.8-9.589.

R Core Team. 2020. *R: A Language and Environment for Statistical Computing*. Manual. Vienna, Austria: R Foundation for Statistical Computing. https://www.R-project.org/.

Rosenberg, Michael S, Julia Koricheva, Jessica Gurevitch, and Kerrie Mengersen. 2013. "Moment and Least-Squares Based Approaches to Meta-Analytic Inference." In *Handbook of Meta-Analysis in Ecology and Evolution*, 108–24. Princeton and Oxford: Princeton University Press.

Shrier, Ian, and Robert W. Platt. 2008. "Reducing Bias Through Directed Acyclic Graphs." *BMC Medical Research Methodology* 8 (1): 70. https://doi.org/10.1186/1471-2288-8-70.

Stan Development Team. 2020. "RStan: The R Interface to Stan." http://mc-stan.org/.

Urli, M., A. J. Porte, H. Cochard, Y. Guengant, R. Burlett, and S. Delzon. 2013. "Xylem Embolism Threshold for Catastrophic Hydraulic Failure in Angiosperm Trees." *Tree Physiology* 33 (7): 672–83. https://doi.org/10.1093/treephys/tpt030.

VanderWeele, Tyler J., Miguel A. Hernán, and James M. Robins. 2008. "Causal Directed Acyclic Graphs and the Direction of Unmeasured Confounding Bias." *Epidemiology (Cambridge, Mass.)* 19 (5): 720–28. https://doi.org/10.1097/EDE.0b013e3181810e29.

Vehtari, Aki, Jonah Gabry, Mans Magnusson, Yuling Yao, and Andrew Gelman. 2019. "Loo: Efficient Leave-One-Out Cross-Validation and WAIC for Bayesian Models." https://mc-stan.org/loo.

Vehtari, Aki, Andrew Gelman, and Jonah Gabry. 2017. "Practical Bayesian Model Evaluation Using Leave-One-Out Cross-Validation and WAIC." *Statistics and Computing* 27 (5): 1413–32. https://doi.org/10.1007/s11222-016-9696-4.

Vehtari, Aki, Andrew Gelman, Daniel Simpson, Bob Carpenter, and Paul-Christian Bürkner. 2020. "Rank-Normalization, Folding, and Localization: An Improved $\widehat{}R{}$ for Assessing Convergence of MCMC." *Bayesian Analysis*. https://doi.org/10.1214/20-BA1221.

Wong, Suan-Chin. 1990. "Elevated Atmospheric Partial-Pressure of $CO_2$ and Plant-Growth. II: Nonstructural Carbohydrate Content in Cotton Plants and Its Effect on Growth-Parameters." *Photosynthesis Research* 23: 171–80.

Yao, Yuling, Aki Vehtari, Daniel Simpson, and Andrew Gelman. 2017. "Using Stacking to Average Bayesian Predictive Distributions." *Bayesian Analysis*. https://doi.org/10.1214/17-BA1091.
