## Supplementary Results for "Mutually inclusive mechanisms of drought-induced tree mortality"

### Supplementary material 2: Supplementary results

Supplementary material for Hajek et al. 2020

#### Supplementary figures

**Figure S2.1** Estimated misclassification rate $\phi$ (posterior mean ± 95% HDI).

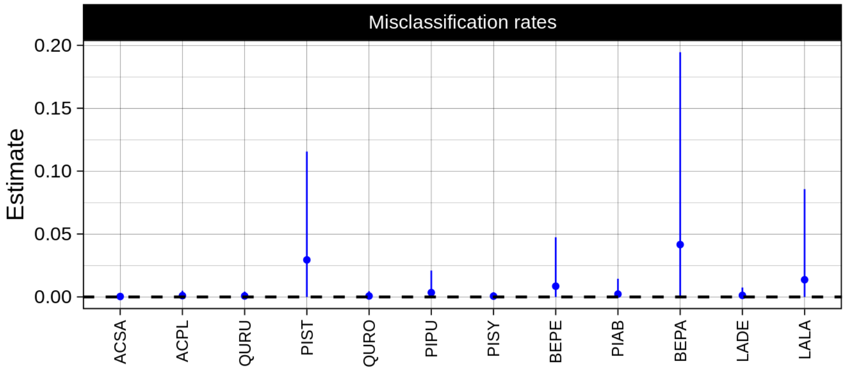


**Figure S2.2** Species level effects of the main predictor variables (on logit scale; posterior mean ± 95% HDI). Blue: credibly reduced risk; red: credibly elevated risk; grey: no credible effect on mortality.

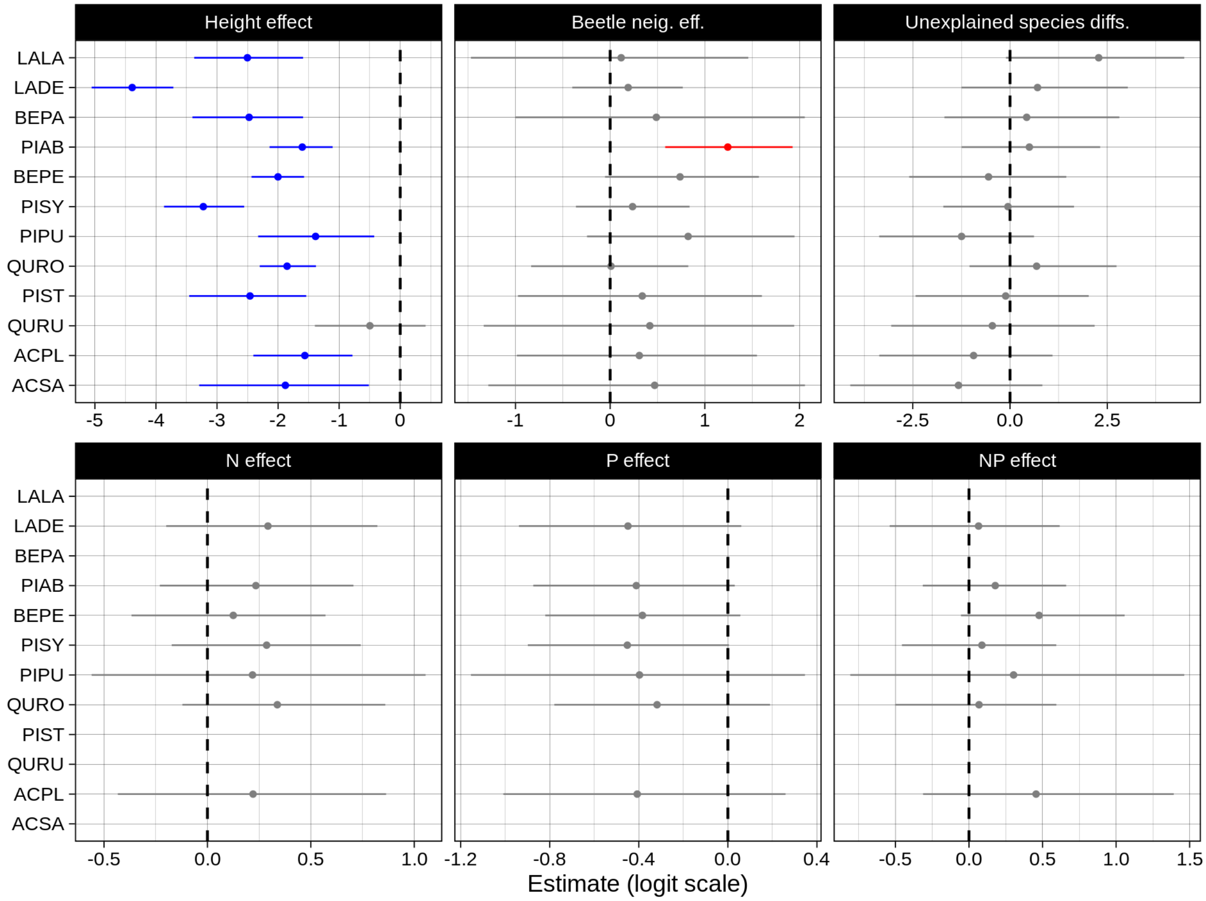

**Figure S2.3** Estimated variance components (on logit scale; posterior mean ± 95% HDI). Left standard deviations of varying effects, and measurement error standard deviation for the NSC measurements. Right: correlation parameters: correlations between the species-wise varying effects parameters, and within-pair correlation for the neighbourhood effects.

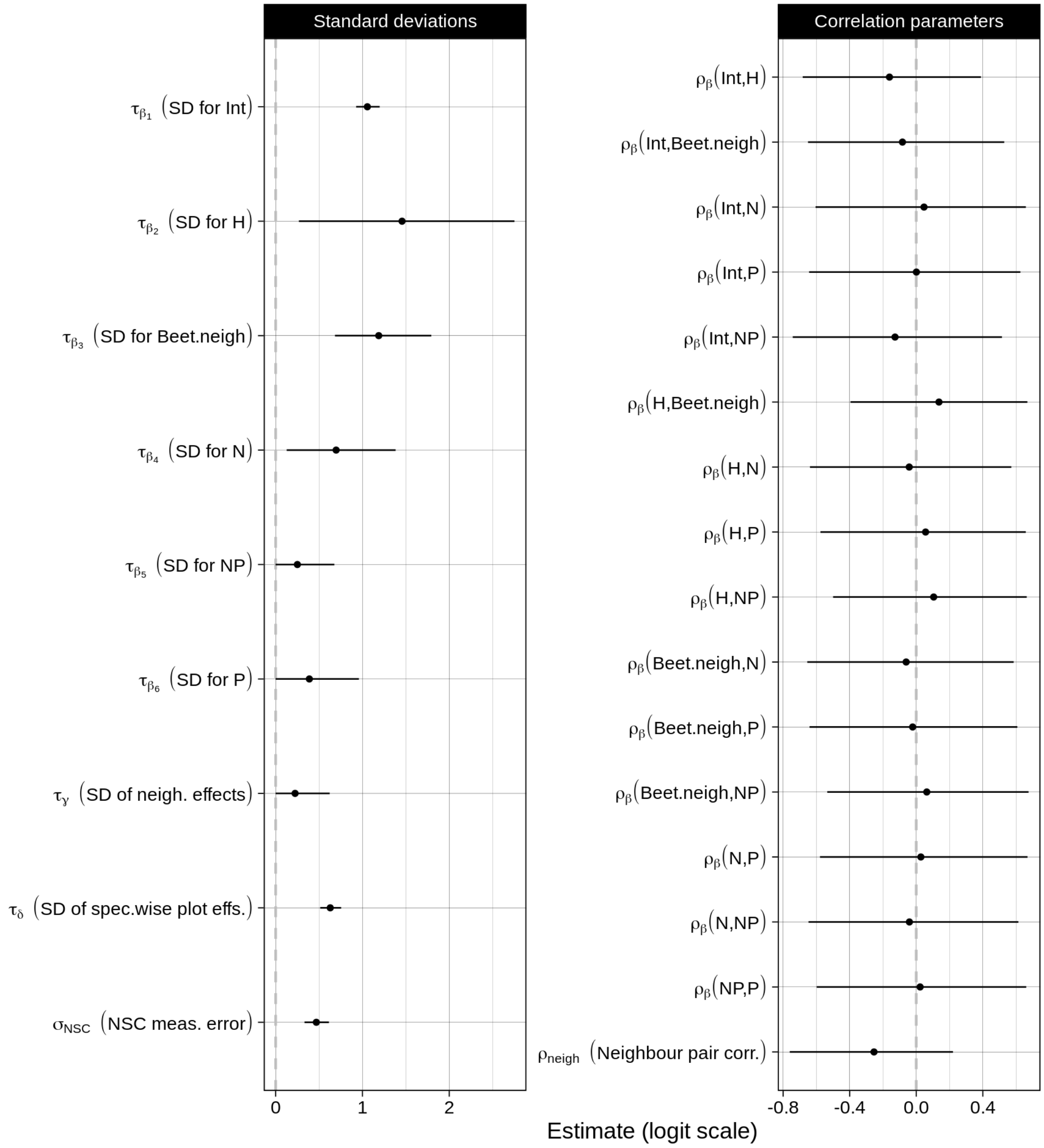

**Figure S2.4** Pairwise directed neighbourhood effect parameters (on logit scale; posterior mean ± 95% HDI) for a) the pairs with the strongest disparities between each partner, b) the pairs with the largest net reduction in mortality and c) the pairs with the largest net increase in predicted mortality. Note that while there are several species combinations that are beneficial for both neighbours, no combination exists that is detrimental for both partners.

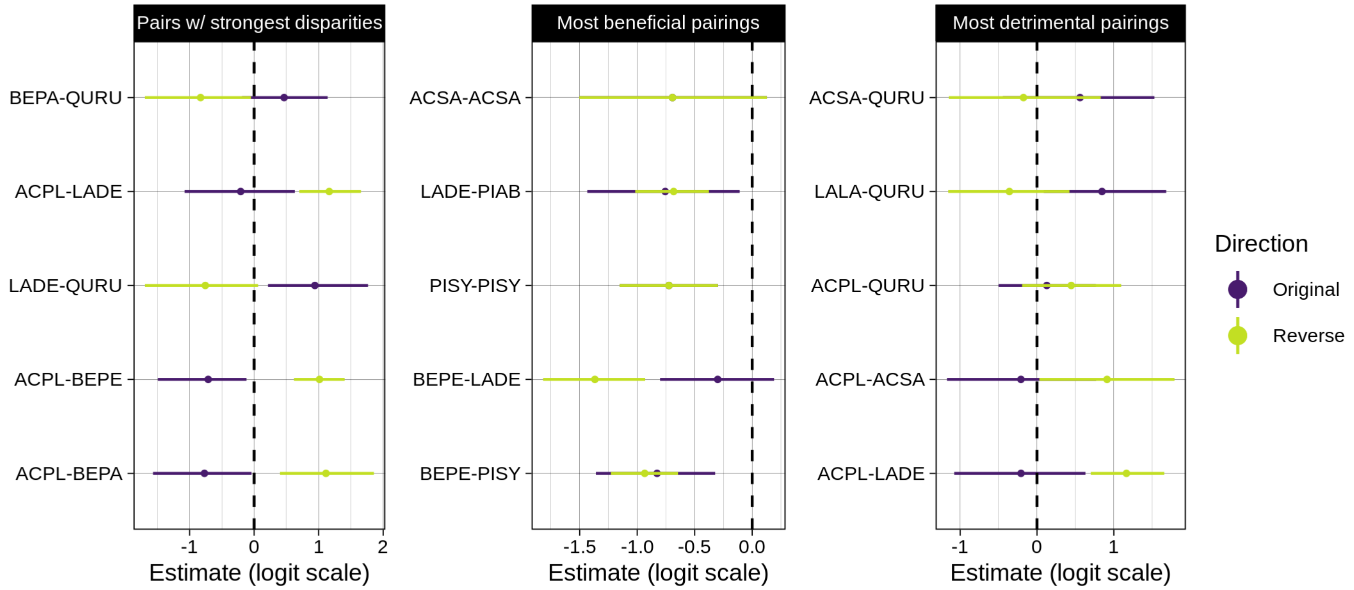


**Figure S2.5** Estimated species-wise plot effects (on logit scale; posterior mean ± 95% HDI) ordered by their average magnitude. Blue: credibly reduced risk; red: credibly elevated risk; grey: no credible effect on mortality.

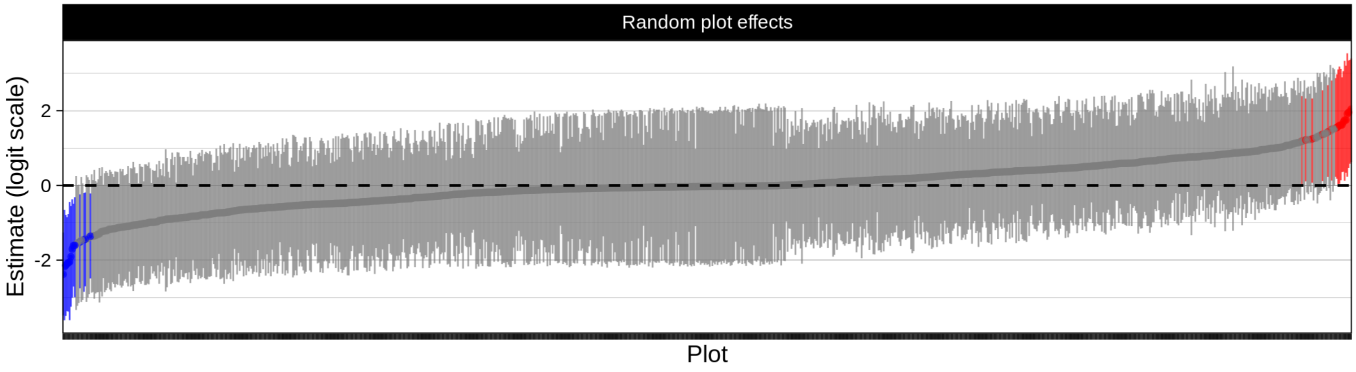

#### Supplementary tables

**Table S2.1** Parameter estimates of the full model. Given are the MCMC mean and standard error (SE), posterior standard deviation (SD), 2.5, 25, 50, 75 and 97.5 % credible intervals, 2.5 and 97.5 % highest posterior density intervals (HDI), effective sample size ($N_{eff}$) and the potential scale reduction factor $\hat{R}$.

| | | | | | Credible intervals | | | | | HDI | | | |
|---|---|---|---|---|---|---|---|---|---|---|---|---|---|
| | Parameter | Mean | SE | SD | 2.5% | 25% | 50% | 75% | 97.5% | 2.5% | 97.5% | $N_{eff}$ | $\hat{R}$ |
| $\beta_0$ | Intercept | -0.558 | 0.007 | 0.567 | -1.701 | -0.923 | -0.541 | -0.183 | 0.550 | -1.747 | 0.500 | 5888.692 | 1.000 |
| | HSM | -1.388 | 0.008 | 0.592 | -2.547 | -1.781 | -1.395 | -1.005 | -0.204 | -2.530 | -0.193 | 5224.623 | 1.000 |
| | Δsugar | 1.931 | 0.011 | 0.715 | 0.436 | 1.499 | 1.949 | 2.411 | 3.276 | 0.492 | 3.311 | 4199.826 | 1.000 |
| | Height | -2.049 | 0.006 | 0.381 | -2.783 | -2.302 | -2.056 | -1.804 | -1.281 | -2.771 | -1.275 | 3686.930 | 1.000 |
| | Beetles | 4.079 | 0.003 | 0.323 | 3.467 | 3.856 | 4.077 | 4.292 | 4.730 | 3.455 | 4.718 | 15884.427 | 1.000 |
| | Beet.n | 0.449 | 0.004 | 0.325 | -0.245 | 0.258 | 0.462 | 0.661 | 1.059 | -0.253 | 1.049 | 6321.805 | 1.000 |
| | N | 0.244 | 0.003 | 0.212 | -0.182 | 0.112 | 0.245 | 0.376 | 0.654 | -0.175 | 0.660 | 6180.332 | 1.000 |
| | NP | 0.230 | 0.003 | 0.253 | -0.269 | 0.078 | 0.226 | 0.383 | 0.738 | -0.264 | 0.740 | 6183.592 | 1.000 |
| | P | -0.395 | 0.003 | 0.206 | -0.797 | -0.525 | -0.395 | -0.266 | 0.016 | -0.801 | 0.009 | 5632.061 | 1.001 |
| $\gamma_0$ | ACPL | 0.535 | 0.003 | 0.206 | 0.142 | 0.396 | 0.528 | 0.668 | 0.953 | 0.120 | 0.928 | 4306.508 | 1.000 |
| | ACSA | -0.035 | 0.003 | 0.224 | -0.468 | -0.184 | -0.036 | 0.113 | 0.413 | -0.479 | 0.399 | 6609.842 | 1.000 |
| | BEPA | -0.531 | 0.003 | 0.220 | -0.973 | -0.675 | -0.525 | -0.381 | -0.112 | -0.964 | -0.103 | 7608.203 | 1.000 |
| | BEPE | -0.746 | 0.003 | 0.191 | -1.139 | -0.874 | -0.741 | -0.615 | -0.382 | -1.148 | -0.392 | 4555.861 | 1.001 |
| | LADE | -0.406 | 0.003 | 0.194 | -0.787 | -0.535 | -0.407 | -0.277 | -0.016 | -0.794 | -0.026 | 4516.691 | 1.001 |
| | LALA | -0.506 | 0.003 | 0.260 | -1.012 | -0.679 | -0.509 | -0.332 | 0.015 | -0.996 | 0.028 | 6175.155 | 1.000 |
| | PIAB | -0.408 | 0.003 | 0.201 | -0.801 | -0.544 | -0.408 | -0.275 | -0.010 | -0.813 | -0.025 | 5965.706 | 1.000 |
| | PIPU | -0.341 | 0.004 | 0.329 | -0.969 | -0.565 | -0.345 | -0.128 | 0.315 | -0.970 | 0.315 | 6388.385 | 1.000 |
| | PIST | -0.350 | 0.003 | 0.261 | -0.870 | -0.521 | -0.351 | -0.178 | 0.162 | -0.889 | 0.138 | 7046.257 | 1.000 |
| | PISY | -0.420 | 0.003 | 0.206 | -0.826 | -0.557 | -0.418 | -0.286 | -0.007 | -0.829 | -0.014 | 5195.302 | 1.002 |
| | QURO | -0.355 | 0.003 | 0.197 | -0.734 | -0.490 | -0.356 | -0.223 | 0.030 | -0.735 | 0.029 | 5989.649 | 1.001 |
| | QURU | 0.416 | 0.003 | 0.251 | -0.058 | 0.246 | 0.409 | 0.580 | 0.926 | -0.066 | 0.916 | 5768.119 | 1.000 |
| c | | 0.486 | 0.001 | 0.060 | 0.365 | 0.447 | 0.489 | 0.527 | 0.596 | 0.371 | 0.600 | 3721.770 | 1.001 |
| $\sigma_{\Delta sugar}$ | | 0.629 | 0.001 | 0.063 | 0.519 | 0.586 | 0.625 | 0.668 | 0.763 | 0.514 | 0.755 | 10410.895 | 1.000 |
| $\tau_\delta$ | | 1.057 | 0.001 | 0.070 | 0.926 | 1.009 | 1.056 | 1.102 | 1.198 | 0.927 | 1.198 | 3428.149 | 1.000 |
| $\tau_\beta$ | Intercept | 1.456 | 0.010 | 0.619 | 0.305 | 1.046 | 1.416 | 1.812 | 2.819 | 0.268 | 2.751 | 3603.415 | 1.001 |
| | Height | 1.187 | 0.004 | 0.301 | 0.734 | 0.975 | 1.144 | 1.348 | 1.902 | 0.683 | 1.793 | 5373.869 | 1.001 |
| | Beet.n | 0.697 | 0.005 | 0.324 | 0.186 | 0.476 | 0.653 | 0.867 | 1.472 | 0.127 | 1.382 | 4775.656 | 1.000 |
| | N | 0.250 | 0.003 | 0.224 | 0.009 | 0.093 | 0.193 | 0.341 | 0.828 | 0.000 | 0.676 | 4508.987 | 1.001 |
| | NP | 0.388 | 0.005 | 0.297 | 0.020 | 0.176 | 0.327 | 0.530 | 1.136 | 0.000 | 0.959 | 3752.965 | 1.001 |
| | P | 0.224 | 0.003 | 0.211 | 0.008 | 0.078 | 0.168 | 0.304 | 0.766 | 0.000 | 0.622 | 5218.982 | 1.000 |
| $\tau_\gamma$ | | 0.469 | 0.001 | 0.073 | 0.344 | 0.417 | 0.463 | 0.514 | 0.630 | 0.333 | 0.614 | 4513.811 | 1.001 |
| $\rho_\gamma$ | | -0.254 | 0.006 | 0.253 | -0.732 | -0.435 | -0.260 | -0.080 | 0.251 | -0.760 | 0.221 | 1637.774 | 1.001 |
| $\rho_\beta$ | Int~H | -0.161 | 0.005 | 0.279 | -0.661 | -0.363 | -0.172 | 0.025 | 0.420 | -0.682 | 0.389 | 3071.307 | 1.001 |
| | Int~Beet.n | -0.083 | 0.003 | 0.313 | -0.655 | -0.312 | -0.091 | 0.137 | 0.527 | -0.650 | 0.529 | 9332.006 | 1.000 |
| | Int~N | 0.047 | 0.003 | 0.332 | -0.599 | -0.191 | 0.051 | 0.292 | 0.668 | -0.605 | 0.659 | 16347.064 | 1.000 |
| | Int~P | 0.001 | 0.003 | 0.338 | -0.638 | -0.246 | -0.001 | 0.244 | 0.637 | -0.644 | 0.626 | 17083.277 | 1.000 |
| | Int~NP | -0.127 | 0.003 | 0.332 | -0.718 | -0.374 | -0.140 | 0.106 | 0.544 | -0.742 | 0.515 | 13816.592 | 1.000 |
| | H~Beet.n | 0.137 | 0.003 | 0.280 | -0.413 | -0.060 | 0.146 | 0.338 | 0.654 | -0.395 | 0.668 | 10929.259 | 1.000 |
| | H~N | -0.042 | 0.003 | 0.321 | -0.634 | -0.281 | -0.044 | 0.188 | 0.578 | -0.638 | 0.571 | 14702.982 | 1.000 |
| | H~P | 0.056 | 0.003 | 0.327 | -0.588 | -0.178 | 0.063 | 0.297 | 0.654 | -0.576 | 0.659 | 14229.812 | 1.000 |
| | H~NP | 0.105 | 0.003 | 0.304 | -0.507 | -0.108 | 0.115 | 0.326 | 0.660 | -0.500 | 0.664 | 12556.964 | 1.000 |
| | Beet.n~N | -0.061 | 0.003 | 0.329 | -0.659 | -0.306 | -0.067 | 0.177 | 0.583 | -0.655 | 0.586 | 8869.477 | 1.000 |
| | Beet.n~P | -0.022 | 0.003 | 0.329 | -0.641 | -0.259 | -0.019 | 0.213 | 0.608 | -0.641 | 0.608 | 11825.554 | 1.000 |
| | Beet.n~NP | 0.063 | 0.003 | 0.322 | -0.556 | -0.169 | 0.066 | 0.300 | 0.657 | -0.535 | 0.675 | 10432.685 | 1.000 |
| | N~P | 0.028 | 0.003 | 0.332 | -0.605 | -0.216 | 0.032 | 0.273 | 0.648 | -0.578 | 0.669 | 9953.418 | 1.000 |
| | N~NP | -0.041 | 0.004 | 0.333 | -0.653 | -0.286 | -0.046 | 0.198 | 0.610 | -0.647 | 0.614 | 8329.151 | 1.000 |
| | NP~P | 0.023 | 0.004 | 0.336 | -0.616 | -0.217 | 0.025 | 0.269 | 0.647 | -0.598 | 0.661 | 8566.358 | 1.000 |

**Table S2.1** (continued).

| Parameter | | Mean | SE | SD | Credible intervals 2.5% | 25% | 50% | 75% | 97.5% | HDI 2.5% | 97.5% | $N_{eff}$ | $\hat{R}$ |
|---|---|---|---|---|---|---|---|---|---|---|---|---|---|
| $\varphi$ | ACPL | 0.001 | 0.000 | 0.002 | 0.000 | 0.000 | 0.000 | 0.001 | 0.006 | 0.000 | 0.005 | 2569.819 | 1.001 |
| | ACSA | 0.000 | 0.000 | 0.001 | 0.000 | 0.000 | 0.000 | 0.000 | 0.003 | 0.000 | 0.002 | 10012.782 | 1.000 |
| | BEPA | 0.042 | 0.001 | 0.067 | 0.000 | 0.000 | 0.002 | 0.064 | 0.225 | 0.000 | 0.195 | 3268.588 | 1.000 |
| | BEPE | 0.009 | 0.000 | 0.017 | 0.000 | 0.000 | 0.000 | 0.008 | 0.060 | 0.000 | 0.047 | 4855.829 | 1.001 |
| | LADE | 0.001 | 0.000 | 0.003 | 0.000 | 0.000 | 0.000 | 0.001 | 0.012 | 0.000 | 0.007 | 9998.864 | 1.000 |
| | LALA | 0.014 | 0.000 | 0.037 | 0.000 | 0.000 | 0.000 | 0.006 | 0.133 | 0.000 | 0.086 | 10367.257 | 1.000 |
| | PIAB | 0.002 | 0.000 | 0.006 | 0.000 | 0.000 | 0.000 | 0.001 | 0.021 | 0.000 | 0.014 | 9627.631 | 1.000 |
| | PIPU | 0.003 | 0.000 | 0.010 | 0.000 | 0.000 | 0.000 | 0.001 | 0.033 | 0.000 | 0.021 | 7806.630 | 1.000 |
| | PIST | 0.029 | 0.001 | 0.041 | 0.000 | 0.000 | 0.005 | 0.053 | 0.132 | 0.000 | 0.116 | 3070.475 | 1.001 |
| | PISY | 0.001 | 0.000 | 0.002 | 0.000 | 0.000 | 0.000 | 0.000 | 0.006 | 0.000 | 0.004 | 10488.031 | 1.000 |
| | QURO | 0.001 | 0.000 | 0.002 | 0.000 | 0.000 | 0.000 | 0.000 | 0.007 | 0.000 | 0.005 | 10229.833 | 1.000 |
| | QURU | 0.001 | 0.000 | 0.002 | 0.000 | 0.000 | 0.000 | 0.000 | 0.007 | 0.000 | 0.004 | 8850.833 | 1.000 |
| $\Delta sugar_{true}$ | ACPL | -0.706 | 0.002 | 0.286 | -1.270 | -0.889 | -0.707 | -0.519 | -0.136 | -1.264 | -0.130 | 14851.140 | 1.000 |
| | ACSA | -0.901 | 0.003 | 0.333 | -1.548 | -1.122 | -0.903 | -0.683 | -0.246 | -1.550 | -0.249 | 15120.181 | 1.000 |
| | BEPA | 0.572 | 0.003 | 0.370 | -0.155 | 0.325 | 0.574 | 0.820 | 1.289 | -0.168 | 1.274 | 14873.211 | 1.000 |
| | BEPE | 1.072 | 0.003 | 0.302 | 0.488 | 0.866 | 1.072 | 1.273 | 1.668 | 0.487 | 1.667 | 10026.191 | 1.000 |
| | LADE | 1.176 | 0.003 | 0.316 | 0.567 | 0.963 | 1.175 | 1.383 | 1.793 | 0.567 | 1.794 | 12235.022 | 1.000 |
| | LALA | 0.481 | 0.004 | 0.315 | -0.137 | 0.267 | 0.477 | 0.693 | 1.100 | -0.125 | 1.109 | 7640.401 | 1.000 |
| | PIAB | -0.443 | 0.003 | 0.291 | -1.022 | -0.638 | -0.441 | -0.247 | 0.126 | -1.029 | 0.113 | 10243.645 | 1.000 |
| | PIPU | -0.207 | 0.003 | 0.294 | -0.801 | -0.402 | -0.203 | -0.014 | 0.374 | -0.815 | 0.352 | 11121.743 | 1.000 |
| | PIST | -1.186 | 0.002 | 0.287 | -1.754 | -1.378 | -1.185 | -0.994 | -0.624 | -1.760 | -0.632 | 14807.185 | 1.000 |
| | PISY | 0.263 | 0.002 | 0.274 | -0.285 | 0.084 | 0.266 | 0.443 | 0.807 | -0.291 | 0.795 | 13659.181 | 1.000 |
| | QURO | 0.470 | 0.003 | 0.284 | -0.094 | 0.281 | 0.472 | 0.661 | 1.025 | -0.090 | 1.028 | 12256.651 | 1.000 |
| | QURU | -0.290 | 0.002 | 0.291 | -0.853 | -0.485 | -0.292 | -0.095 | 0.278 | -0.845 | 0.283 | 17797.151 | 1.000 |
| $HSM_{true}$ | ACPL | 1.121 | 0.000 | 0.066 | 0.993 | 1.077 | 1.121 | 1.165 | 1.253 | 0.999 | 1.258 | 20830.459 | 1.000 |
| | ACSA | 0.038 | 0.002 | 0.282 | -0.516 | -0.150 | 0.038 | 0.228 | 0.580 | -0.526 | 0.567 | 20263.579 | 1.000 |
| | BEPA | -1.345 | 0.001 | 0.198 | -1.736 | -1.478 | -1.344 | -1.211 | -0.966 | -1.723 | -0.957 | 21889.075 | 1.000 |
| | BEPE | -0.926 | 0.000 | 0.054 | -1.031 | -0.963 | -0.927 | -0.889 | -0.822 | -1.032 | -0.823 | 21428.029 | 1.000 |
| | LADE | -0.200 | 0.001 | 0.113 | -0.422 | -0.276 | -0.199 | -0.125 | 0.025 | -0.435 | 0.010 | 22537.909 | 1.000 |
| | LALA | -0.463 | 0.000 | 0.057 | -0.577 | -0.502 | -0.463 | -0.425 | -0.353 | -0.573 | -0.350 | 24648.544 | 1.000 |
| | PIAB | -0.244 | 0.003 | 0.334 | -0.891 | -0.470 | -0.242 | -0.017 | 0.412 | -0.880 | 0.418 | 15209.480 | 1.000 |
| | PIPU | 0.312 | 0.000 | 0.052 | 0.211 | 0.277 | 0.311 | 0.346 | 0.414 | 0.209 | 0.410 | 20705.774 | 1.000 |
| | PIST | -0.705 | 0.000 | 0.063 | -0.829 | -0.748 | -0.705 | -0.663 | -0.581 | -0.827 | -0.580 | 23367.241 | 1.000 |
| | PISY | -0.635 | 0.001 | 0.134 | -0.896 | -0.725 | -0.635 | -0.544 | -0.371 | -0.895 | -0.369 | 21778.931 | 1.000 |
| | QURO | 0.970 | 0.001 | 0.111 | 0.752 | 0.894 | 0.970 | 1.044 | 1.187 | 0.746 | 1.181 | 19505.473 | 1.000 |
| | QURU | 1.894 | 0.002 | 0.328 | 1.258 | 1.671 | 1.891 | 2.119 | 2.526 | 1.257 | 2.524 | 17442.808 | 1.000 |
| $\beta_{Int}$ | ACPL | -0.937 | 0.016 | 1.139 | -3.483 | -1.626 | -0.813 | -0.135 | 1.022 | -3.362 | 1.092 | 5388.917 | 1.000 |
| | ACSA | -1.325 | 0.017 | 1.288 | -4.199 | -2.117 | -1.176 | -0.385 | 0.782 | -4.106 | 0.830 | 5675.225 | 1.000 |
| | BEPA | 0.428 | 0.013 | 1.118 | -1.693 | -0.285 | 0.357 | 1.106 | 2.806 | -1.686 | 2.809 | 7084.369 | 1.000 |
| | BEPE | -0.553 | 0.013 | 0.996 | -2.559 | -1.154 | -0.521 | 0.037 | 1.484 | -2.592 | 1.447 | 6150.985 | 1.000 |
| | LADE | 0.707 | 0.016 | 1.094 | -1.282 | -0.023 | 0.634 | 1.395 | 3.010 | -1.247 | 3.028 | 4492.298 | 1.000 |
| | LALA | 2.279 | 0.020 | 1.236 | 0.004 | 1.392 | 2.265 | 3.122 | 4.740 | -0.104 | 4.478 | 3872.866 | 1.000 |
| | PIAB | 0.497 | 0.011 | 0.889 | -1.236 | -0.071 | 0.454 | 1.060 | 2.333 | -1.243 | 2.317 | 6699.399 | 1.000 |
| | PIPU | -1.246 | 0.014 | 1.050 | -3.456 | -1.927 | -1.177 | -0.471 | 0.547 | -3.363 | 0.616 | 5899.133 | 1.001 |
| | PIST | -0.112 | 0.013 | 1.092 | -2.445 | -0.745 | -0.060 | 0.572 | 2.008 | -2.427 | 2.022 | 6898.929 | 1.000 |
| | PISY | -0.053 | 0.010 | 0.833 | -1.751 | -0.574 | -0.042 | 0.457 | 1.614 | -1.714 | 1.644 | 6425.015 | 1.000 |
| | QURO | 0.683 | 0.013 | 0.951 | -1.119 | 0.045 | 0.629 | 1.284 | 2.680 | -1.042 | 2.738 | 5491.528 | 1.000 |
| | QURU | -0.454 | 0.016 | 1.297 | -3.199 | -1.214 | -0.369 | 0.331 | 2.056 | -3.054 | 2.175 | 6260.304 | 1.000 |

**Table S2.1** (continued).

| Parameter | | Mean | SE | SD | Credible intervals 2.5% | 25% | 50% | 75% | 97.5% | HDI 2.5% | 97.5% | $N_{eff}$ | $\hat{R}$ |
|---|---|---|---|---|---|---|---|---|---|---|---|---|---|
| $\beta_{Height}$ | ACPL | 0.488 | 0.007 | 0.496 | -0.508 | 0.160 | 0.492 | 0.823 | 1.444 | -0.511 | 1.440 | 5290.378 | 1.000 |
| | ACSA | 0.169 | 0.007 | 0.716 | -1.240 | -0.295 | 0.170 | 0.642 | 1.567 | -1.241 | 1.567 | 10017.690 | 1.000 |
| | BEPA | -0.425 | 0.007 | 0.548 | -1.579 | -0.787 | -0.404 | -0.054 | 0.596 | -1.534 | 0.625 | 5478.473 | 1.000 |
| | BEPE | 0.048 | 0.006 | 0.409 | -0.789 | -0.218 | 0.057 | 0.323 | 0.833 | -0.742 | 0.865 | 4271.599 | 1.000 |
| | LADE | -2.340 | 0.007 | 0.472 | -3.320 | -2.643 | -2.328 | -2.009 | -1.462 | -3.280 | -1.431 | 4963.505 | 1.000 |
| | LALA | -0.453 | 0.007 | 0.541 | -1.538 | -0.807 | -0.446 | -0.089 | 0.577 | -1.547 | 0.564 | 5573.104 | 1.001 |
| | PIAB | 0.446 | 0.006 | 0.417 | -0.397 | 0.172 | 0.452 | 0.721 | 1.257 | -0.360 | 1.284 | 4559.720 | 1.000 |
| | PIPU | 0.663 | 0.007 | 0.551 | -0.421 | 0.293 | 0.663 | 1.029 | 1.763 | -0.417 | 1.769 | 6946.401 | 1.001 |
| | PIST | -0.410 | 0.008 | 0.554 | -1.599 | -0.755 | -0.390 | -0.039 | 0.620 | -1.555 | 0.658 | 5452.946 | 1.000 |
| | PISY | -1.175 | 0.006 | 0.442 | -2.106 | -1.457 | -1.161 | -0.869 | -0.342 | -2.033 | -0.293 | 4738.814 | 1.001 |
| | QURO | 0.197 | 0.006 | 0.397 | -0.618 | -0.059 | 0.205 | 0.458 | 0.966 | -0.598 | 0.974 | 4394.963 | 1.000 |
| | QURU | 1.553 | 0.006 | 0.535 | 0.512 | 1.196 | 1.555 | 1.902 | 2.626 | 0.533 | 2.640 | 6780.730 | 1.000 |
| $\beta_{Beet.n}$ | ACPL | -0.141 | 0.005 | 0.590 | -1.414 | -0.478 | -0.094 | 0.220 | 0.976 | -1.344 | 1.027 | 13440.895 | 1.000 |
| | ACSA | 0.021 | 0.006 | 0.752 | -1.533 | -0.374 | 0.012 | 0.426 | 1.567 | -1.521 | 1.574 | 14296.805 | 1.000 |
| | BEPA | 0.039 | 0.006 | 0.699 | -1.297 | -0.351 | 0.007 | 0.401 | 1.549 | -1.340 | 1.504 | 14299.984 | 1.000 |
| | BEPE | 0.289 | 0.005 | 0.460 | -0.551 | -0.013 | 0.254 | 0.563 | 1.275 | -0.544 | 1.277 | 8759.421 | 1.000 |
| | LADE | -0.259 | 0.004 | 0.403 | -1.050 | -0.522 | -0.260 | -0.006 | 0.569 | -1.093 | 0.513 | 8176.416 | 1.000 |
| | LALA | -0.333 | 0.007 | 0.697 | -1.892 | -0.726 | -0.280 | 0.105 | 0.945 | -1.827 | 0.992 | 11087.427 | 1.000 |
| | PIAB | 0.793 | 0.006 | 0.445 | 0.013 | 0.483 | 0.765 | 1.075 | 1.742 | -0.035 | 1.650 | 4964.792 | 1.000 |
| | PIPU | 0.374 | 0.006 | 0.559 | -0.636 | 0.004 | 0.319 | 0.709 | 1.614 | -0.692 | 1.530 | 9045.731 | 1.000 |
| | PIST | -0.110 | 0.005 | 0.613 | -1.428 | -0.448 | -0.089 | 0.240 | 1.104 | -1.378 | 1.137 | 14971.748 | 1.000 |
| | PISY | -0.213 | 0.004 | 0.400 | -1.019 | -0.464 | -0.207 | 0.027 | 0.604 | -1.046 | 0.574 | 8071.496 | 1.000 |
| | QURO | -0.440 | 0.005 | 0.455 | -1.442 | -0.723 | -0.405 | -0.123 | 0.374 | -1.386 | 0.392 | 7326.669 | 1.000 |
| | QURU | -0.030 | 0.007 | 0.745 | -1.660 | -0.421 | 0.001 | 0.406 | 1.395 | -1.611 | 1.440 | 12531.943 | 1.000 |
| $\beta_{N}$ | ACPL | -0.024 | 0.002 | 0.259 | -0.613 | -0.116 | -0.006 | 0.078 | 0.522 | -0.574 | 0.556 | 10881.780 | 1.000 |
| | ACSA | -0.014 | 0.003 | 0.345 | -0.720 | -0.118 | -0.003 | 0.097 | 0.675 | -0.725 | 0.666 | 10074.482 | 1.000 |
| | BEPA | 0.001 | 0.003 | 0.315 | -0.671 | -0.099 | 0.001 | 0.105 | 0.684 | -0.618 | 0.725 | 10650.454 | 1.000 |
| | BEPE | -0.119 | 0.003 | 0.223 | -0.673 | -0.220 | -0.061 | 0.009 | 0.214 | -0.651 | 0.230 | 5209.746 | 1.001 |
| | LADE | 0.048 | 0.003 | 0.250 | -0.445 | -0.065 | 0.017 | 0.160 | 0.625 | -0.455 | 0.612 | 8055.078 | 1.000 |
| | LALA | 0.028 | 0.004 | 0.359 | -0.689 | -0.095 | 0.007 | 0.147 | 0.814 | -0.770 | 0.715 | 10087.265 | 1.000 |
| | PIAB | -0.010 | 0.002 | 0.215 | -0.486 | -0.100 | -0.003 | 0.082 | 0.445 | -0.485 | 0.445 | 7947.255 | 1.000 |
| | PIPU | -0.026 | 0.003 | 0.345 | -0.760 | -0.130 | -0.006 | 0.082 | 0.651 | -0.731 | 0.666 | 11671.330 | 1.000 |
| | PIST | 0.006 | 0.003 | 0.311 | -0.619 | -0.094 | 0.001 | 0.102 | 0.642 | -0.613 | 0.648 | 10424.718 | 1.000 |
| | PISY | 0.042 | 0.003 | 0.216 | -0.382 | -0.055 | 0.015 | 0.130 | 0.556 | -0.422 | 0.510 | 7073.813 | 1.000 |
| | QURO | 0.094 | 0.003 | 0.223 | -0.268 | -0.022 | 0.042 | 0.186 | 0.658 | -0.293 | 0.624 | 7163.232 | 1.000 |
| | QURU | -0.012 | 0.003 | 0.344 | -0.765 | -0.122 | -0.002 | 0.103 | 0.687 | -0.716 | 0.736 | 10570.540 | 1.000 |
| $\beta_{NP}$ | ACPL | 0.226 | 0.005 | 0.405 | -0.369 | -0.009 | 0.120 | 0.397 | 1.269 | -0.404 | 1.188 | 5723.784 | 1.000 |
| | ACSA | 0.052 | 0.005 | 0.475 | -0.903 | -0.130 | 0.017 | 0.228 | 1.094 | -0.866 | 1.125 | 10015.894 | 1.000 |
| | BEPA | -0.026 | 0.004 | 0.451 | -0.997 | -0.198 | -0.009 | 0.146 | 0.915 | -0.963 | 0.946 | 11307.691 | 1.000 |
| | BEPE | 0.246 | 0.004 | 0.307 | -0.208 | 0.024 | 0.182 | 0.417 | 0.976 | -0.227 | 0.948 | 4977.782 | 1.001 |
| | LADE | -0.165 | 0.004 | 0.322 | -0.895 | -0.339 | -0.112 | 0.023 | 0.411 | -0.850 | 0.450 | 7209.952 | 1.000 |
| | LALA | -0.103 | 0.005 | 0.518 | -1.257 | -0.302 | -0.046 | 0.108 | 0.895 | -1.194 | 0.952 | 9970.717 | 1.000 |
| | PIAB | -0.052 | 0.004 | 0.282 | -0.685 | -0.188 | -0.022 | 0.096 | 0.484 | -0.667 | 0.498 | 6477.171 | 1.000 |
| | PIPU | 0.073 | 0.004 | 0.472 | -0.889 | -0.113 | 0.030 | 0.255 | 1.128 | -0.844 | 1.157 | 11111.740 | 1.000 |
| | PIST | -0.009 | 0.004 | 0.450 | -0.970 | -0.180 | -0.005 | 0.163 | 0.932 | -0.970 | 0.932 | 11565.159 | 1.000 |
| | PISY | -0.142 | 0.004 | 0.292 | -0.814 | -0.293 | -0.089 | 0.021 | 0.376 | -0.783 | 0.401 | 6286.326 | 1.000 |
| | QURO | -0.162 | 0.004 | 0.294 | -0.849 | -0.313 | -0.104 | 0.013 | 0.325 | -0.823 | 0.345 | 6339.599 | 1.000 |
| | QURU | 0.060 | 0.005 | 0.519 | -0.973 | -0.139 | 0.023 | 0.256 | 1.191 | -1.004 | 1.140 | 10746.521 | 1.000 |

**Table S2.1** (continued).

| Parameter | | Mean | SE | SD | Credible intervals 2.5% | 25% | 50% | 75% | 97.5% | HDI 2.5% | 97.5% | $N_{eff}$ | $\hat{R}$ |
|---|---|---|---|---|---|---|---|---|---|---|---|---|---|
| $\beta_P$ | ACPL | -0.012 | 0.003 | 0.251 | -0.565 | -0.090 | 0.000 | 0.079 | 0.492 | -0.559 | 0.497 | 9254.688 | 1.000 |
| | ACSA | -0.002 | 0.003 | 0.301 | -0.635 | -0.092 | 0.000 | 0.090 | 0.615 | -0.624 | 0.620 | 8192.272 | 1.000 |
| | BEPA | -0.009 | 0.003 | 0.278 | -0.593 | -0.096 | -0.002 | 0.078 | 0.556 | -0.595 | 0.549 | 8609.692 | 1.000 |
| | BEPE | 0.012 | 0.002 | 0.196 | -0.397 | -0.066 | 0.003 | 0.087 | 0.442 | -0.391 | 0.445 | 6755.533 | 1.001 |
| | LADE | -0.054 | 0.003 | 0.231 | -0.629 | -0.144 | -0.017 | 0.054 | 0.374 | -0.581 | 0.408 | 6913.104 | 1.001 |
| | LALA | 0.007 | 0.003 | 0.331 | -0.663 | -0.096 | 0.001 | 0.106 | 0.704 | -0.663 | 0.704 | 8939.855 | 1.000 |
| | PIAB | -0.016 | 0.003 | 0.204 | -0.488 | -0.096 | -0.004 | 0.070 | 0.419 | -0.441 | 0.455 | 6432.249 | 1.001 |
| | PIPU | -0.002 | 0.003 | 0.310 | -0.682 | -0.092 | 0.001 | 0.093 | 0.626 | -0.691 | 0.609 | 8917.516 | 1.000 |
| | PIST | -0.012 | 0.003 | 0.276 | -0.612 | -0.096 | -0.002 | 0.075 | 0.543 | -0.630 | 0.522 | 8572.779 | 1.000 |
| | PISY | -0.056 | 0.003 | 0.207 | -0.572 | -0.135 | -0.020 | 0.039 | 0.309 | -0.540 | 0.334 | 6737.108 | 1.000 |
| | QURO | 0.077 | 0.002 | 0.208 | -0.262 | -0.027 | 0.028 | 0.154 | 0.613 | -0.290 | 0.578 | 7022.809 | 1.000 |
| | QURU | 0.029 | 0.003 | 0.325 | -0.600 | -0.076 | 0.007 | 0.121 | 0.753 | -0.627 | 0.724 | 8740.867 | 1.000 |
